# Adaptive evolution shapes the present-day distribution of the thermal sensitivity of population growth rate

**DOI:** 10.1101/712885

**Authors:** Dimitrios - Georgios Kontopoulos, Thomas P. Smith, Timothy G. Barraclough, Samraat Pawar

**Affiliations:** Science and Solutions for a Changing Planet DTP, Imperial College London, London, United Kingdom; Department of Life Sciences, Imperial College London, Silwood Park, Ascot, Berkshire, United Kingdom

## Abstract

Developing a thorough understanding of how ectotherm physiology adapts to different thermal environments is of crucial importance, especially in the face of global climate change. A key aspect of an organism’s thermal performance curve—the relationship between fitness-related trait performance and temperature—is its thermal sensitivity, i.e., the rate at which trait values increase with temperature within its typically-experienced thermal range. For a given trait, the distribution of thermal sensitivities across species, often quantified as “activation energy” values, is typically right-skewed. Currently, the mechanisms that generate this distribution are unclear, with considerable debate about the role of thermodynamic constraints vs adaptive evolution. Here, using a phylogenetic comparative approach, we study the evolution of the thermal sensitivity of population growth rate across phytoplankton (Cyanobacteria and eukaryotic microalgae) and prokaryotes (bacteria and archaea), two microbial groups that play a major role in the global carbon cycle. We find that thermal sensitivity across these groups is moderately phylogenetically heritable, and that its distribution is shaped by repeated evolutionary convergence throughout its parameter space. More precisely, we detect bursts of adaptive evolution in thermal sensitivity, increasing the amount of overlap among its distributions in different clades. We obtain qualitatively similar results from evolutionary analyses of the thermal sensitivities of two physiological rates underlying growth rate: net photosynthesis and respiration of plants. Furthermore, we find that these episodes of evolutionary convergence are consistent with two opposing forces: decrease in thermal sensitivity due to environmental fluctuations and increase due to adaptation to stable environments. Overall, our results indicate that adaptation can lead to large and relatively rapid shifts in thermal sensitivity, especially in microbes where rapid evolution can occur at short time scales. Thus, more attention needs to be paid to elucidating the implications of rapid evolution in organismal thermal sensitivity for ecosystem functioning.

**Author summary:** Changes in environmental temperature influence the performance of biological traits (e.g., respiration rate) in ectotherms, with the relationship between trait performance and temperature (the “thermal performance curve”) being single-peaked. Understanding how thermal performance curves adapt to different environments is important for predicting how organisms will be impacted by climate change. One key aspect of the shape of these curves is the thermal sensitivity near temperatures typically experienced by the species. Whether and how thermal sensitivity responds to different environments is a topic of active debate. To shed light on this, here we perform an evolutionary analysis of the thermal sensitivity of three key traits of prokaryotes, phytoplankton, and plants. We show that thermal sensitivity does not evolve in a gradual manner, but can differ considerably even between closely related species. This suggests that thermal sensitivity undergoes rapid adaptive evolution, which is further supported by our finding that thermal sensitivity varies weakly with latitude. We conclude that variation in thermal sensitivity arises partly from adaptation to environmental factors and that this may need to be accounted for in ecophysiological models.

## Introduction

Current climate change projections suggest that the average global temperature in 2100 will be higher than the average of 1986-2005 by 0.3-4.8°C [1], coupled with an increase in temperature fluctuations in certain areas [2]. Therefore, it is now more important than ever to understand how temperature changes affect biological systems, from individuals to whole ecosystems. At the level of individual organisms, temperature affects functional traits in the form of the “thermal performance curve” (TPC). Typically, this TPC, especially when the trait is a rate (e.g., respiration rate, photosynthesis, growth), takes the shape of a negatively-skewed unimodal curve (Fig. 1) [3,4]. The curve increases (approximately) exponentially to a maximum (*T*_pk_), and then also decreases exponentially, with the fall being steeper than the rise. Understanding how various aspects of the shape of this TPC adapt to a changing thermal environment is crucial for predicting how rapidly organisms can respond to climate change.

**Fig 1.**
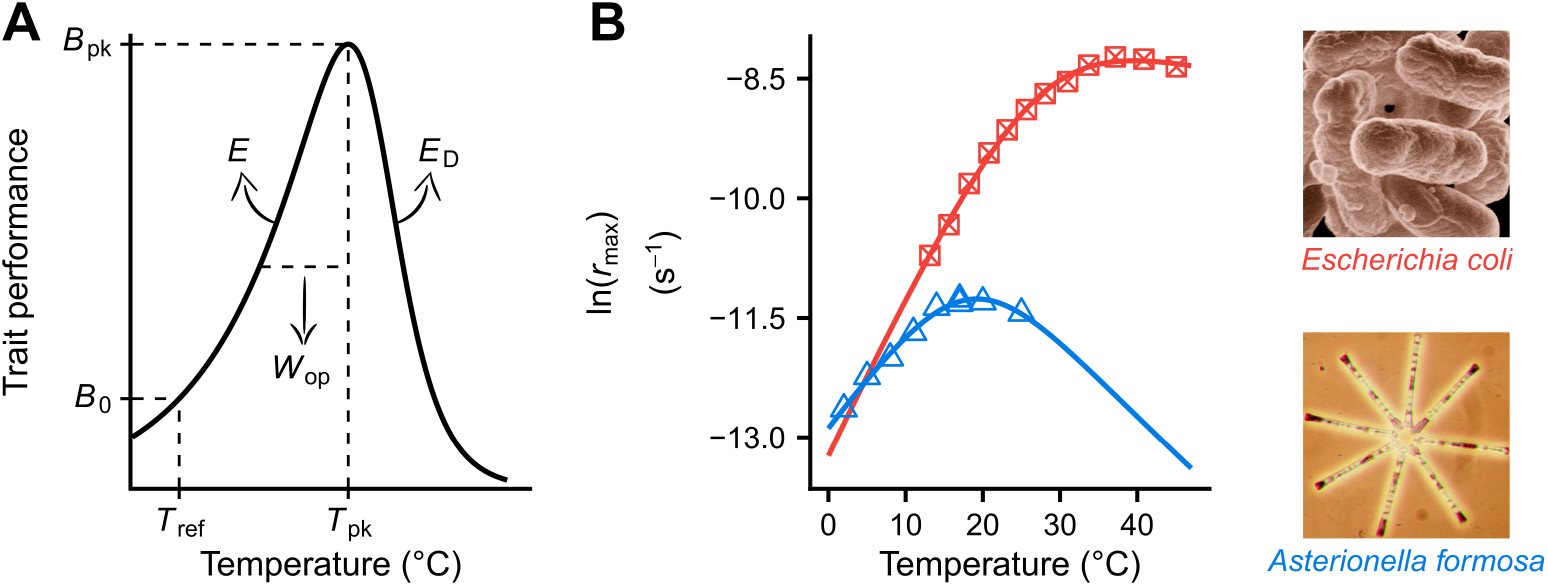
The thermal performance curve (TPC) of ectotherm metabolic traits, as described by the Sharpe-Schoolfield model [5]. (A) *T*_pk_ (K) is the temperature at which the curve peaks, reaching a maximum height that is equal to *B*_pk_ (in units of trait performance). *E* and *E*_D_ (eV) control how smoothly the TPC rises and falls respectively. *B*_0_ (in units of trait performance) is the trait performance normalised at a reference temperature (*T*_ref_) below the peak. In addition, *W*_op_ (K), the operational niche width of the TPC, can also be calculated *a posteriori* as the difference between *T*_pk_ and the temperature at the rise of the TPC where *B*(*T*) = 0.5 · *B*_pk_. This niche width metric assumes that species routinely experience temperatures well below *T*_pk_, consistent with the results of previous studies [6, 7]. (B) TPCs of individual- and population-level traits (such as *r*_max_) are usually well-described by the Sharpe-Schoolfield model.

According to the Metabolic Theory of Ecology (MTE) as well as a large body of physiological research, the shape of the TPC is expected to reflect the effects of temperature on the kinetics of a single rate-limiting enzyme involved in key metabolic reactions [5, 8–11]. Under this assumption, the rise in trait values up to *T*_pk_ can be mechanistically described using the Boltzmann-Arrhenius equation:

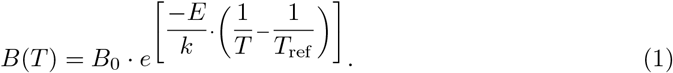

Here, *B* is the value of a biological trait, *B*_0_ is a normalisation constant—that includes the effect of cell or body size—which gives the trait value at a reference temperature (*T*_ref_), *T* is temperature (in K), *k* is the Boltzmann constant (8.617 · 10^−5^ eV · K^−1^), and *E* (eV) is the thermal sensitivity of the trait at the rising component of the TPC up to *T*_pk_. Because *T*_pk_ tends to be higher than the mean environmental temperature [6, 7, 12], *E* represents the thermal sensitivity within the organism’s typically-experienced thermal range.

Early MTE studies argued that, because of strong thermodynamic constraints, adaptation will predominantly result in changes in *B*_0_, whereas *E* will remain almost constant across traits (e.g., respiration rate, *r*_max_), species, and environments around a range of 0.6-0.7 eV [8–10]. The latter assumption is referred to in the literature as “universal temperature dependence” (UTD). This restricted range of values that *E* can take is centered on the putative mean activation energy of respiration (≈ 0.65 eV). A notable exception to the UTD is photosynthesis rate, which is expected to have a lower *E* value of ≈ 0.32 eV, reflecting the activation energy of the rate-limiting steps of photosynthesis [13].

The existence of a UTD has been strongly debated. From a theoretical standpoint, critics of the UTD have argued that the Boltzmann-Arrhenius model is too simple to mechanistically describe the complex physiological mechanisms of diverse organisms [3,14–16], and is inadequate for describing TPCs emerging from the interaction of multiple factors, and not just the effects of temperature on enzyme kinetics. That is, the *E* calculated by fitting the Boltzmann-Arrhenius model to biological traits is an emergent property that does not directly reflect the activation energy of a single rate-limiting enzyme. For example, a fixed thermal sensitivity for net photosynthesis rate is not realistic because it depends on the rate of gross photosynthesis as well as photorespiration, which is in turn determined not only by temperature but also by the availability of CO_2_ in relation to O_2_ [17].

Indeed, there is now overwhelming empirical evidence for variation in *E* (thermal sensitivity) far exceeding the narrow 0.6-0.7 eV range, with such variation being, to an extent, taxonomically-structured [12,18–23]. Furthermore, the distribution of *E* values across species is typically not Gaussian but right-skewed. If we assume that *E* is nearly constant across species and, therefore, variation in *E* is mainly due to measurement error, such skewness could be the outcome of the proximity of the *E* distribution to its lower boundary (0 eV). In that case, however, we would expect a high density of *E* values close to 0 eV, but such a pattern has not been observed [18]. Both the deviations from the MTE expectation of a heavily restricted range for *E* and the shape of its distribution have been argued to be partly driven by adaptation to local environmental factors by multiple studies. These include selection on prey to have lower thermal sensitivity than predators (the “thermal life-dinner principle”) [18], adaptation to temperature fluctuations within and/or across generations [3,21,24–26], and adaptive increases in carbon allocation or use efficiency due to warming [27–30].

In general then, adaptive changes in the TPCs of underlying (fitness-related) traits are expected to influence the TPCs of higher-order traits such as *r*_max_, resulting in deviations from a UTD. Therefore, understanding how the thermal sensitivity of *r*_max_ and its distribution evolves is particularly important, as it may also yield useful insights about the evolution of the TPCs of underlying physiological traits (e.g., respiration rate, photosynthesis rate, and carbon allocation efficiency). Indeed, systematic shifts in the thermal sensitivity of fundamental physiological traits have been documented [27,31–33], albeit not through comparative analyses of large datasets.

In particular, phylogenetic heritability—the extent to which closely related species have more similar trait values than species chosen at random—can provide key insights regarding the evolution of thermal sensitivity. A phylogenetic heritability of 1 indicates that the evolution of the trait across the phylogenetic tree is indistinguishable from a random walk (Brownian motion) in the parameter space. Note that this does not necessarily indicate that the trait evolves neutrally, as it may be under selection towards a non-stationary optimum that itself performs a random walk [34]. In contrast, a phylogenetic heritability of 0 indicates that trait values are independent of the phylogeny. This is the case either because i) the trait is practically invariant across species and any variation is due to measurement error, or ii) the evolution of the trait is very fast and with frequent convergence (i.e., independent evolution of similar trait values by different lineages). It is worth clarifying that rapid trait evolution that does not result in convergence (e.g., when major clades are extremely separated in the parameter space) will not lead to a complete absence of phylogenetic heritability. Phylogenetic heritabilities between 0 and 1 reflect deviations from Brownian motion (e.g., due to occasional patterns of evolutionary convergence). Among phytoplankton, measures of thermal sensitivity of *r*_max_ (*E* and *W*_op_) have previously been shown to exhibit intermediate phylogenetic heritability [35]. This indicates that among phytoplankton, thermal sensitivity is not constant but evolves along the phylogeny, albeit not as a purely random walk in trait space, reflecting either thermodynamically constrained evolution or rapid evolution in response to selection.

To understand i) how variation in thermal sensitivity accumulates across multiple autotroph and heterotroph groups, and ii) whether its distribution is shaped by environmental selection, here we conduct a thorough investigation of the evolutionary patterns of thermal sensitivity, focusing particularly on *r*_max_. Using a phylogenetic comparative approach, we test the following hypotheses:

1. **Thermal sensitivity does not evolve across species and any variation is noise-like.** In this scenario, thermodynamic constraints would force *E* to be distributed around a mean of 0.65 eV (or 0.32 eV in the case of photosynthesis), with deviations from the mean being mostly due to measurement error. Depending on the magnitude of the error, the *E* distribution would either be approximately Gaussian (little measurement error) or non-Gaussian with a high density near 0 eV (substantial measurement error). This hypothesis agrees with the UTD concept of early MTE studies. If this hypothesis holds, thermal sensitivity would have zero phylogenetic heritability and would not vary systematically across different environments.
2. **Thermal sensitivity evolves gradually across species but tends to revert to a key central value, without ever moving very far from it.** This hypothesis is also consistent with the UTD assumption, as it is a relaxed variant of hypothesis 1. Here, small deviations from the central tendency of 0.65 eV (or 0.32 eV) are possible, as they would reflect adaptation of species’ enzymes to certain ecological lifestyles or niches. Therefore, thermal sensitivity would be weakly phylogenetically heritable. Thermodynamic constraints would prevent large deviations from the central tendency.
3. **Thermal sensitivity evolves in other ways.** This is an “umbrella” hypothesis that encompasses multiple sub-hypotheses that do not invoke the UTD assumption. For example, a key central tendency (thermodynamic constraint) may still exist, but its influence would be very weak, allowing for a wide exploration of the parameter space away from it. In this case, changes in thermal sensitivity could be the outcome of adaptation to different thermal environments. Another sub-hypothesis is that clades differ systematically in the rate at which thermal sensitivity evolves, due to the occasional emergence of evolutionary innovations. Thus, clades with high evolutionary rates would be able to better explore the parameter space of thermal sensitivity (i.e., through large changes in *E* and *W*_op_ values), compared to low-rate clades in which thermal sensitivity would evolve more gradually. A third sub-hypothesis is that evolution may favour species (and metabolic variants) that are relatively insensitive to temperature fluctuations. In that case, the central tendency of *E* would not be stationary, but moving towards lower values with evolutionary time. It is worth clarifying that these three sub-hypotheses are not necessarily mutually exclusive.

## Results

### Dataset sources

We combined two pre-existing datasets of *r*_max_ TPCs, spanning 380 phytoplankton species (a polyphyletic group that includes prokaryotic Cyanobacteria and eukaryotic phyla such as Dinophyta) [35] and 272 prokaryote species (bacteria and archaea) [32]. In addition, we also collected two TPC datasets of traits that underlie *r*_max_: net photosynthesis and respiration rates of algae, aquatic and terrestrial plants (221 and 201 species respectively) [30]. We used these two smaller datasets to understand if the evolutionary patterns of thermal sensitivity differ between i) higher-order traits and ii) traits that are more tightly linked to organismal physiology. Trait values were typically measured under nutrient-, light- and CO_2_-saturated conditions (where applicable), after acclimation to each experimental temperature.

To investigate the evolution of measures of thermal sensitivity across species, we reconstructed the phylogeny of as many species in the four datasets as possible, from publicly available nucleotide sequences of i) the small subunit rRNA gene from all species groups and the ii) cbbL/rbcL gene from photosynthetic prokaryotes, algae, and plants (see the Methods section). We managed to obtain small subunit rRNA gene sequences from 537 species and cbbL/rbcL sequences from 208 of them (Tables S4 and S5 in the S1 Appendix).

TPC parameters were quantified for each species/strain present in the phylogeny using the Sharpe-Schoolfield model (see Fig. 1 and the Methods section). The resulting estimates of *E* (the slope of the rise of the TPC) and *W*_op_ (the operational niche width of the TPC) were found to be right-skewed (Fig. S2 in Appendix S1) as has been shown previously [18,21]. Furthermore, we did not detect a disproportionately high density of thermal sensitivity values near the lower boundary of *E* (0 eV), as we would expect if all variation was due to strong measurement error around a true value of e.g., 0.65 eV. Thus, these results are not consistent with the hypothesis of a nearly invariant thermal sensitivity (hypothesis 1).

### Phylogenetic comparative analyses

We next investigated the evolutionary patterns of thermal sensitivity. Given that the main focus of this study is to investigate how the thermal sensitivity of *r*_max_ (a direct measure of fitness) evolves, most of the following comparative analyses were performed on our two large TPC datasets (*r*_max_ of phytoplankton and prokaryotes). Besides this, the sample sizes of the two smaller datasets would be inadequate for obtaining robust results for many of our analyses. If an analysis makes use of all four datasets, this is explicitly stated.

An issue that is worth mentioning is the overlap between the datasets of phytoplankton and prokaryotic TPCs, given that both of them include Cyanobacteria. To address this, we kept Cyanobacteria as part of the phytoplankton dataset (due to their functional similarity) and did not include them in analyses of prokaryotes. We also examined whether our results were mainly driven by the long evolutionary distance between Cyanobacteria and eukaryotic phytoplankton by repeating all phytoplankton analyses after removing Cyanobacteria (see subsection S3.2 in Appendix S1).

### Estimation of phylogenetic heritability

As TPC parameters capture different features of the shape of the same curve, it is likely that some of them may covary [35]. To account for this in the estimation of phylogenetic heritability, we fitted a multi-response phylogenetic regression model using the MCMCglmm R package (v. 2.26) [36] in which all TPC parameters formed a combined response. To compare the phylogenetic heritabilities of TPC parameters between planktonic photosynthetic autotrophs and other microbes (autotrophs and heterotrophs), we fitted the model separately to our two large TPC datasets: *r*_max_ of phytoplankton and prokaryotes. To satisfy the assumption of models of trait evolution that the change in trait values is normally distributed, we transformed all TPC parameters so that their distributions would be approximately Gaussian (see Fig. 2).

**Fig 2.**
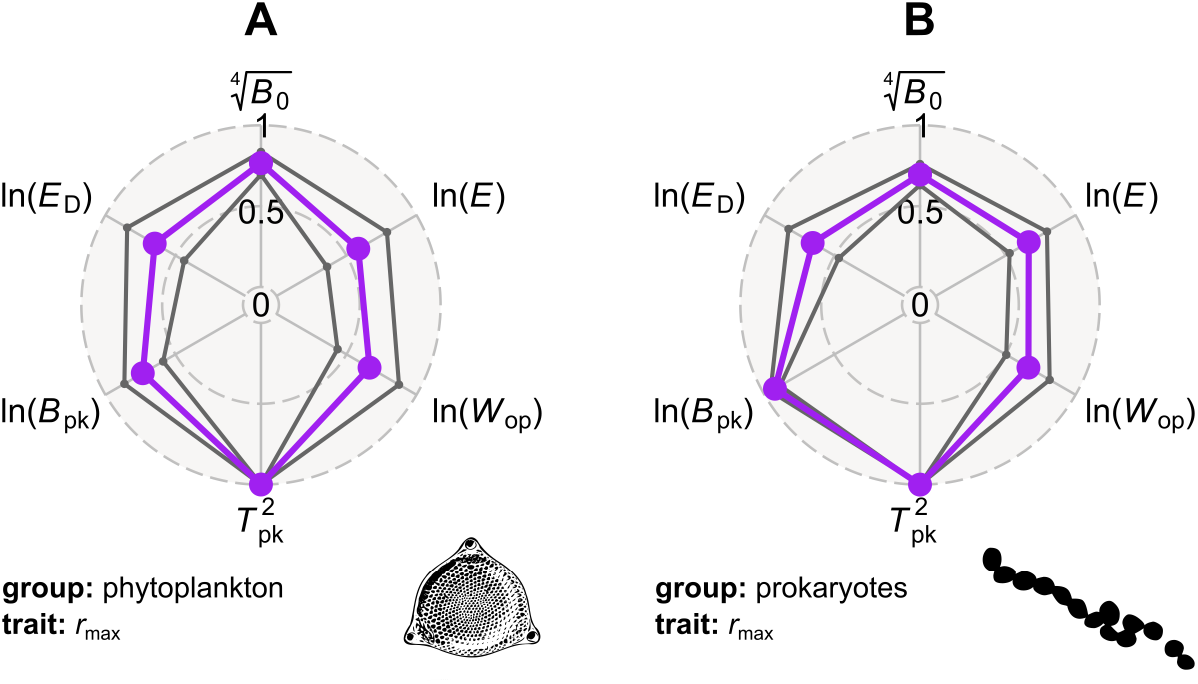
Moderate to strong phylogenetic heritability can be detected in all TPC parameters, across phytoplankton and prokaryotes. The three circles of each radar chart correspond to phylogenetic heritabilities of 0, 0.5, and 1. Mean phylogenetic heritability estimates are shown in purple, whereas the 95% HPD intervals are in dark grey. Note that we transformed all TPC parameters so that their statistical distributions would be approximately Gaussian.

Non-negligible phylogenetic heritability was detected in measures of thermal sensitivity (*E* and *W*_op_), as well as all other TPC parameters, across phytoplankton (including or excluding Cyanobacteria) and prokaryotes (Figs. 2 and S8). In particular, the phylogenetic heritability estimates of ln(*E*) and ln(*W*_op_) were statistically different from both zero and one, indicating that the two TPC parameters evolve across the phylogeny but not in a purely random (Brownian motion) manner. It is worth stressing that even the lower bounds of the 95% highest posterior density (HPD) intervals of ln(*E*) and ln(*W*_op_) were far greater than zero, allowing us to completely rule out the possibility that all variation in thermal sensitivity is due to measurement error. In general, TPC parameters exhibit a similar phylogenetic heritability between the two species groups. The only major exception is ln(*B*_pk_), which is considerably more heritable among prokaryotes than among phytoplankton. This difference in phylogenetic heritability most likely reflects the strength of the positive correlation between *B*_pk_ and *T*_pk_ (a “hotter is better” pattern) in the two groups. More precisely, *T*_pk_, which has a phylogenetic heritability of ≈ 1, is more strongly correlated with *B*_pk_ among prokaryotes [32] than among phytoplankton [35], possibly due to the differences in their cellular physiology. For example, phytoplankton growth rate depends on the interplay among the processes of photosynthesis, respiration, as well as cell maintenance, whose thermal sensitivities can strongly differ [30]. Overall, these results serve as further evidence that hypothesis 1 (that thermal sensitivity does not vary across species) can clearly be rejected.

### Partitioning of thermal sensitivity across the phylogeny

To understand why thermal sensitivity has an intermediate phylogenetic heritability, we examined how clades throughout the phylogeny explore the parameter space (of *E* and *W*_op_) using a disparity-through-time analysis [37,38]. At each branching point of the phylogeny, mean subclade disparity is calculated as the average squared Euclidean distance among trait values within the subclades, normalised to the disparity of trait values across the entire tree. Mean subclade disparity values close to 0 indicate that the average trait variance within subclades is much lower than that across the entire phylogeny. When the opposite occurs, the mean subclade disparity will be close to 1 or even higher. The resulting disparity line is then compared to the null expectation, i.e., an envelope of disparities obtained from simulations of Brownian motion on the same tree. Through the comparison of the true trait disparity with the null expectation, it is possible to identify the periods of evolutionary time during which mean subclade disparity is higher or lower than expected under Brownian motion. Higher than expected subclade disparity indicates that clades converge in trait space, whereas lower than expected subclade disparity indicates that clades occupy distinct areas of parameter space. The latter pattern is consistent with an adaptive radiation, in which an initial period of rapid trait evolution is typically followed by a deceleration of the evolutionary rate as ecological niches become filled [39,40]. Frequent episodes of higher than expected subclade disparity (evolutionary convergence) in thermal sensitivity or segregation of major clades in the parameter space would be consistent with hypothesis 3.

The mean subclade disparity of thermal sensitivity measures was considerably higher than expected near the present, highlighting an increasing overlap in the parameter space of thermal sensitivity among distinct clades (Figs. 3 and S9). This pattern of increasing clade-wide convergence in thermal sensitivity is also apparent when comparing the thermal sensitivity distributions of different phyla (Figs. 4 and S3). For example, *E* and *W*_op_ are similarly distributed among Proteobacteria and Bacillariophyta despite the long evolutionary distance that separates them. This high convergence in thermal sensitivity space by diverse lineages suggests that variation in the two TPC parameters is mainly driven by adaptation to local environmental conditions, irrespective of species’ evolutionary history. In other words, it is likely that particular thermal strategies (e.g., having low thermal sensitivity) may yield significant fitness gains in certain environments (e.g., those with strong temperature fluctuations that occur predominantly across—rather than within—generations [24,25]), leading to convergent evolution of thermal sensitivity. It is worth noting that these disparity patterns are not an artefact of a potentially inaccurate tree topology, as higher than expected subclade disparity occurs mainly near the present, where tree nodes have generally high statistical support (Fig. S1).

**Fig 3.**
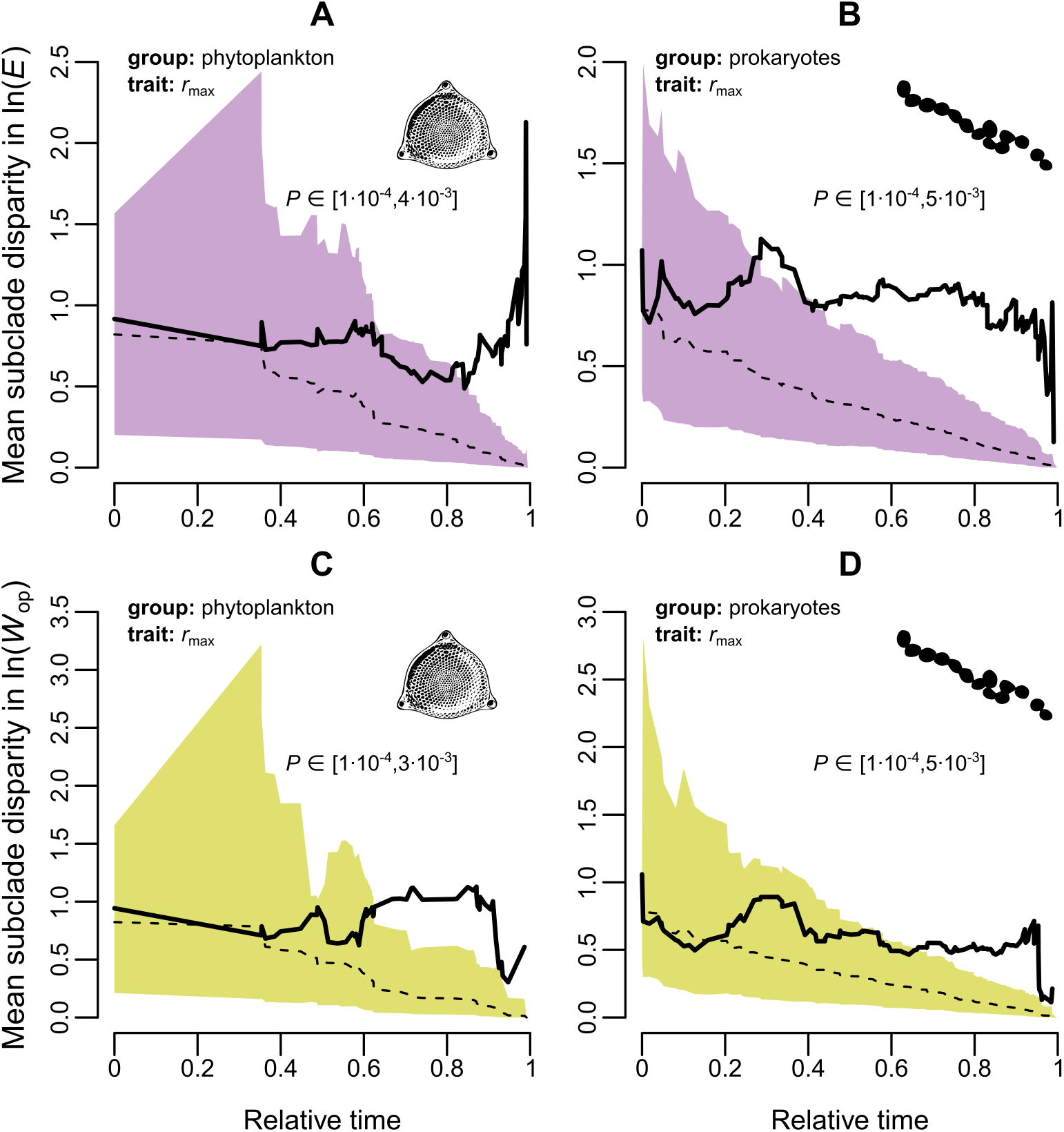
Change in mean subclade disparity in thermal sensitivity through time. Shaded regions represent the 95% confidence interval of the resulting trait disparity from 10,000 simulations of random Brownian evolution on each respective subtree (subset of the entire phylogeny). The dashed line stands for the median disparity across simulations, whereas the solid line is the observed trait disparity. The latter is plotted from the root of the tree (*t* = 0) until the most recent internal node. The reported *P*-values were obtained from the rank envelope test [38], whose null hypothesis is that the trait follows a random walk in the parameter space. Note that instead of a single value, a range of *P*-values is produced for each panel, due to the existence of ties. In general, species from evolutionarily remote clades tend to increasingly overlap in thermal sensitivity space (mean subclade disparity exceeds that expected under Brownian motion) with time.

**Fig 4.**
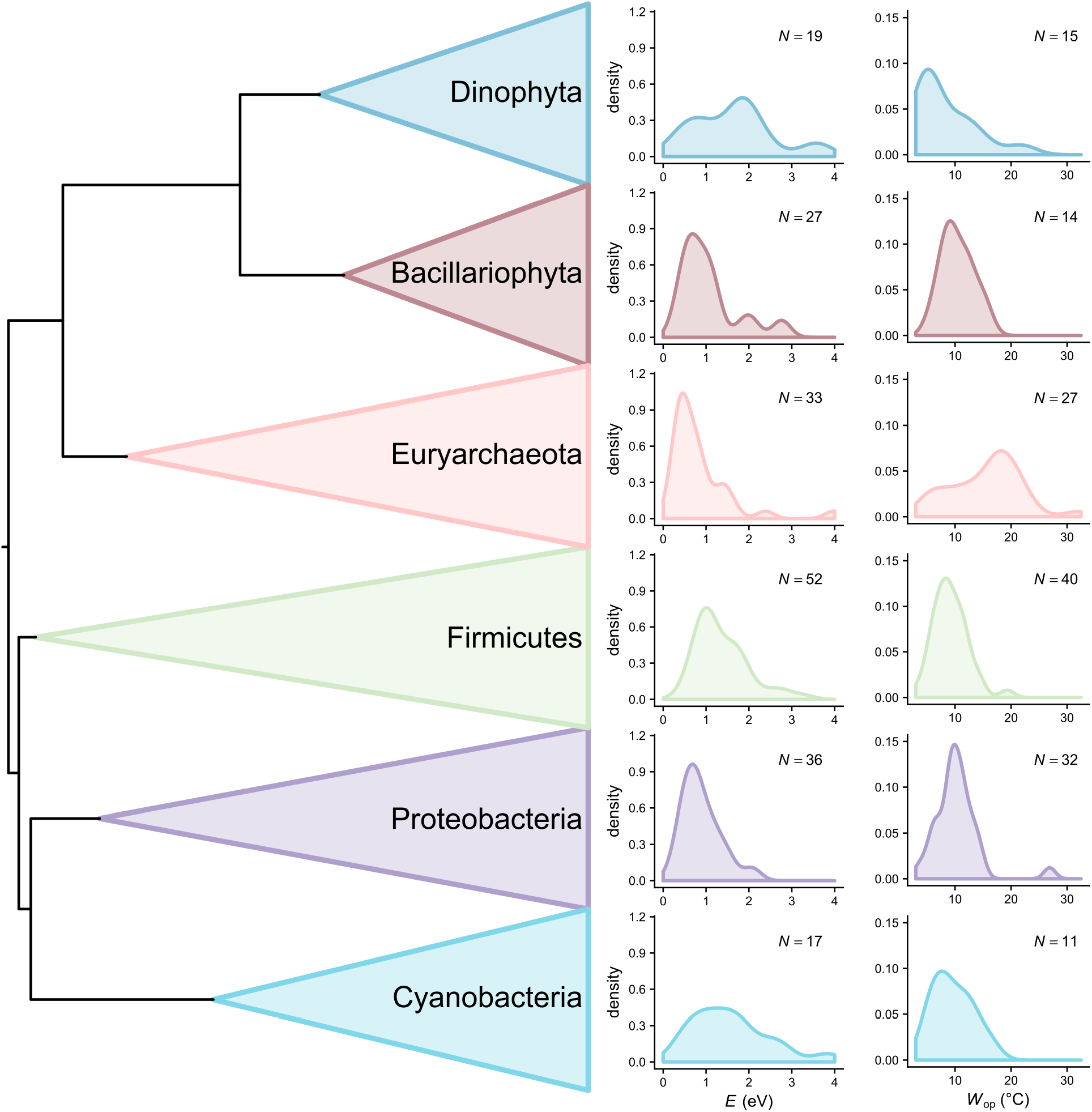
Distributions of thermal sensitivity estimates of *r*_max_ for the largest (most species-rich) phyla of this study. In general, more variation can be observed within than among phyla.

### Mapping the evolutionary rate on the phylogeny

We next investigated if clades systematically differ in their evolutionary rate for thermal sensitivity (part of hypothesis 3). To this end, we examined the variation in the evolutionary rate of thermal sensitivity measures across the phylogeny by fitting three extensions of the Brownian motion model: the free model [41], the stable model [42], and the Lévy model [43]. Under the free model, the trait takes a random walk in the parameter space (Brownian motion) but with an evolutionary rate that varies across branches. The stable model can be seen as a generalisation of the free model, as the evolutionary change in trait values is sampled from a heavy-tailed stable distribution, of which the Gaussian distribution (assumed under Brownian motion) is a special case. Thus, the stable model should provide a more accurate representation of evolutionary rate variation, as it is better able to accommodate jumps in parameter space towards rare and extreme trait values. Finally, the Lévy model represents evolution under Brownian motion combined with occasional episodes of rapid trait change.

The results were robust to the choice of model used for inferring evolutionary rates (Figs. 5,S5, and S6). Rate shifts tend to occur sporadically throughout the phylogeny and especially in late-branching lineages, without being limited to particular clades. This pattern suggests that there is little systematic variation in the evolutionary rate of thermal sensitivity among clades, with sudden bursts of trait evolution arising in parallel across evolutionarily remote lineages.

**Fig 5.**
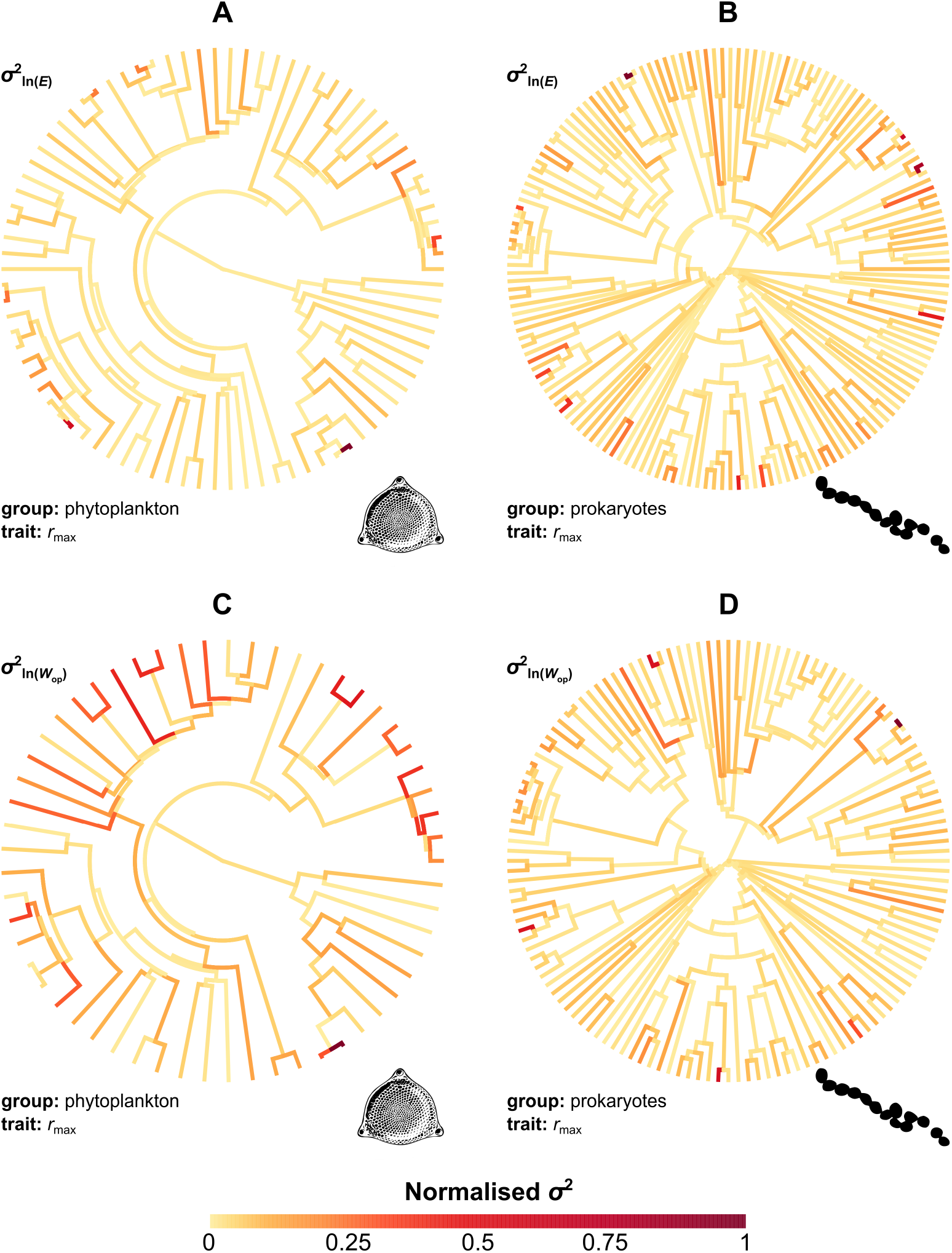
Variation in the evolutionary rate of thermal sensitivity across the phylogeny. Rates were estimated by fitting the stable model of trait evolution to each dataset and were then normalised between 0 and 1. Most branches exhibit relatively low rates of evolution (orange), whereas the highest rates (red and brown) are generally observed in late-branching lineages across different clades.

### Visualization of trait evolution as a function of time, and test for directional selection

To further describe the evolution of thermal sensitivity, we visualized the *E* and *W*_op_ values from the root of each subtree until the present day, across all four TPC datasets. Ancestral states – and the uncertainty around them – were obtained from fits of the stable model of trait evolution, as described in the previous subsection. The visualization allowed us to test hypothesis 2, i.e., that thermal sensitivity evolves closely around a central value of 0.65 eV (or 0.32 eV), with large deviations from this value reverting quickly back to it. To this end, and to also test the hypothesis of directional selection towards lower thermal sensitivity (part of hypothesis 3), we used the following model:

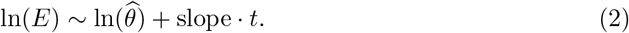

ln(*E*) values (those from extant species and ancestral states inferred with the stable model) were regressed against a central value (ln(*θ*)) and a slope that captures a putative linear trend towards lower/higher values with relative time, *t*. The same model was also fitted to ln(*W*_op_). The regressions were performed with MCMCglmm and were corrected for phylogeny as this resulted in lower Deviance Information Criterion (DIC) [44] values than those obtained from non-phylogenetic variants of the models. More precisely, we executed two MCMCglmm chains per regression for a million generations, sampling every thousand generations after the first hundred thousand.

This analysis (Fig. 6) did not provide support for the hypothesis of strongly constrained adaptive evolution around a single key central value (hypothesis 2). Instead, lineages explore large parts of the parameter space, often moving rapidly towards the upper and lower bounds (i.e., 0 and 4 eV), without reverting back to the presumed central tendency (e.g., see the clade denoted by the arrow in Fig. 6D). The estimated central values for *E* of the two *r*_max_ datasets were much higher than the MTE expectation and, in the case of prokaryotes (Fig. 6B), the 95% HPD interval did not include 0.65. Similarly, the inferred central values for *E* of net photosynthesis rate and respiration rate (0.52 eV and 2.06 eV respectively; Fig. S10A,B) were both higher than 0.32 and 0.65 eV. The slope parameter that would capture the effects of directional selection in thermal sensitivity (part of hypothesis 3) was not statistically different from zero for any dataset.

**Fig 6.**
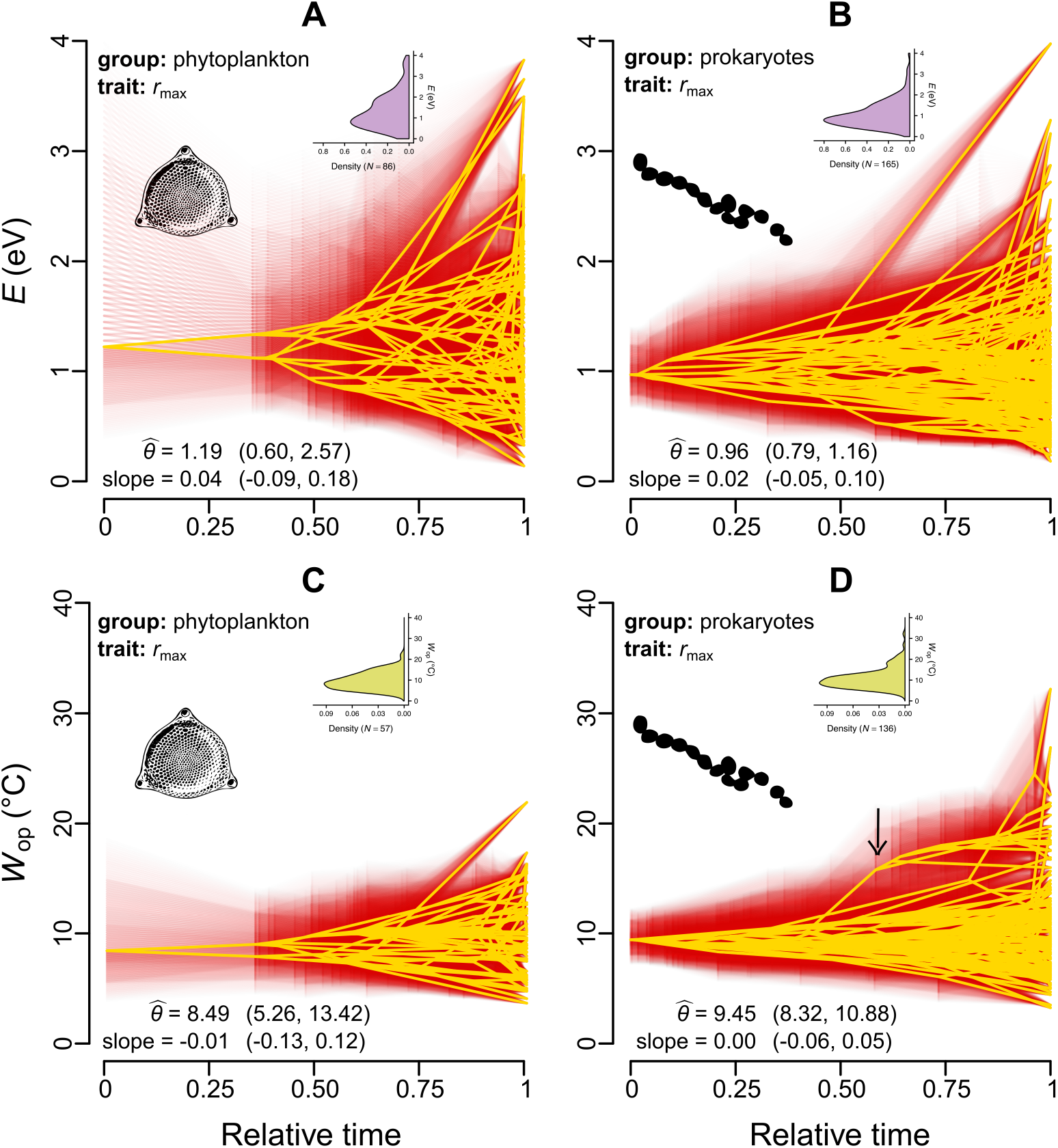
Projection of the phylogeny into thermal sensitivity versus time space. The values of ancestral nodes were estimated from fits of the stable model. Yellow lines represent the median estimates, whereas the 95% credible intervals are shown in red. 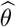 is the estimated central tendency for each panel, whereas the existence of a linear trend towards lower/higher values is captured by the reported slope. Parentheses stand for the 95% HPD intervals for 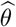 and the slope. All estimates were obtained for ln(*E*) and ln(*W*_op_), but the parameters are shown here in linear scale. The inset figures show the density distributions of *E* and *W*_op_ values of extant species in the dataset. The arrow in panel D shows an example of a whole clade shifting towards high *W*_op_ values, without being attracted back to 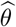.

### Latitudinally structured variation in thermal sensitivity

All our analyses so far converge on one conclusion: that the evolution of the thermal sensitivities of fitness-related traits can be rapid and largely independent of the evolutionary history of each lineage. This suggests that certain environments may select for particular values of thermal sensitivity. To identify environmental adaptation in thermal sensitivity, we tested for latitudinal variation in it using the combination of all four TPC datasets. Specifically, the increase in temperature fluctuations from low to intermediate absolute latitudes is expected to increasingly select for thermal generalists (lower *E* and higher *W*_op_ values) [3,45,46]. At high latitudes, however, temperature fluctuations may further increase or progressively decrease, depending on environment type (marine versus terrestrial) and differences between the two hemispheres [3,45,46]. In any case, the overwhelming majority of our thermal sensitivity estimates belonged to species/strains from low and intermediate latitudes (Fig. S11), enabling us to investigate the hypothesized gradual transition towards lower thermal sensitivity from the equator to intermediate latitudes.

Latitude indeed explained a significant amount of variation in *E* (which declined as expected) but not in *W*_op_ (Figs. 7 and S12, Tables S1 and S2). The *E* estimates of *r*_max_, net photosynthesis rate, and respiration rate differed statistically in their intercepts but not in their slopes against latitude, although the latter could be an artefact of the small sample size. This result suggests that latitude could influence the *E* values of not only *r*_max_ but also other traits across various species groups. Dividing latitude into three bins (i.e., low, intermediate, and high absolute latitudes) and comparing their *E* distributions yielded similar conclusions (Fig. S13, Table S3).

**Fig 7.**
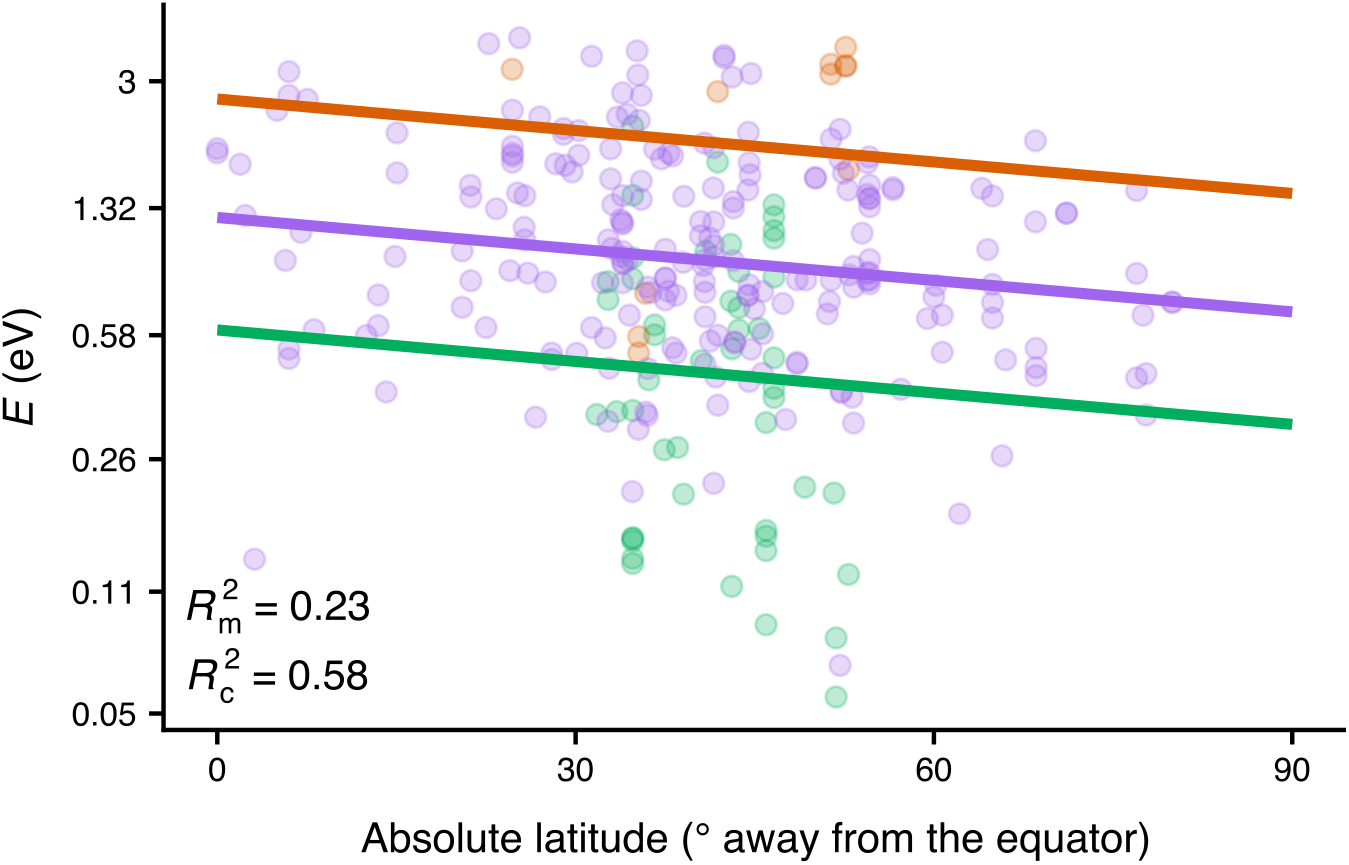
*E* values weakly decrease with absolute latitude. 23% of the variance is explained by latitude and trait identity, which increases to 58% if species identity is added as a random effect on the intercept. Note that values on the vertical axis increase exponentially.

We also tested for a possible latitudinal clade age bias which could arise if certain clades originated in particular latitudes and only much later expanded to other areas [47,48]. For this, we performed a Mantel test [49] to estimate the correlation between phylogenetic distance and latitudinal distance for the two largest groups of our study (phytoplankton and prokaryotes). No such bias was detected for phytoplankton (*r* = 0.04, *P* = 0.114), whereas, for prokaryotes, the correlation was statistically supported but very weak (*r* = 0.11, *P* = 0.002). This result indicates that neither species group is characterised by very strong dispersal limitation throughout its evolutionary history.

## Discussion

In this study, we have performed a thorough analysis of the evolution of the thermal sensitivities of *r*_max_ in phytoplankton and prokaryotes and its two key underlying physiological traits in plants (net photosynthesis rate and respiration rate). To achieve this, we formulated and tested three alternative hypotheses that represent different views expressed in the literature regarding the impact of thermodynamic constraints on the evolution of thermal sensitivity of fitness-related traits.

The first hypothesis was that the activity of a single key rate-limiting enzyme of respiration or photosynthesis directly determines the performance of physiological traits [9,13] and emergent proxies for fitness (such as *r*_max_) [50,51] (the Universal Temperature Dependence assumption). As a result, thermal sensitivity should be strictly constant across traits, species, and environments. This hypothesis was first introduced in early papers that described the Metabolic Theory of Ecology [8–10]. In contrast to the UTD expectation, we detected substantial variation in thermal sensitivity, within and across traits and species groups (Fig. S2 in the S1 Appendix). Furthermore, the distribution of *E* (slope of the rising part of the TPC) values did not exhibit an inflated density near its lower boundary (around 0 eV), as we would expect if all variation in thermal sensitivity was due to measurement error. The rejection of hypothesis 1 was additionally supported by our finding that thermal sensitivity is moderately phylogenetically heritable across phytoplankton and prokaryotes.

Our second hypothesis was that thermal sensitivity evolves across species, but remains close to a key value imposed by strong (but not insurmountable) thermodynamic constraints. We tested this hypothesis using a series of phylogenetic comparative analyses which revealed that the evolution of thermal sensitivity is characterised by an increasing overlap in parameter space by evolutionarily remote lineages (Figs. 3 and 4) due to bursts of rapid evolution (Fig. 5). Additionally, visualisation of thermal sensitivity evolution through time (Figs. 6 and S10) showed that thermal sensitivity can rapidly move away from its presumed central value without being strongly attracted back to it (e.g., see the arrow in Fig. 6D). In conclusion, these results lead us to reject hypothesis 2, i.e., that thermal sensitivity evolves under very strong thermodynamic constraints.

Our final hypothesis was that thermal sensitivity evolves in an adaptive manner and that even if a central tendency exists, its influence on thermal sensitivity evolution is weak. This hypothesis was supported by the results of all phylogenetic comparative analyses and by our detection of a systematic relationship between *E* and latitude. The latter is likely driven by the increase in temperature fluctuations from the equator to intermediate latitudes and agrees with the expectation that thermally-variable environments should select for phenotypes with low thermal sensitivity, and vice versa [3,21,24,25]. That a similar latitudinal effect could not be detected on *W*_op_ (the operational niche width of the TPC) is possibly because of the much smaller sample size available for this, combined with the fact that *W*_op_ is nonlinearly related to *E* (Fig. S4). In any case, *E* is arguably a more meaningful measure of niche width than *W*_op_ because the latter assumes that species mainly experience temperatures close to *T*_pk_, while *E* captures the entire rise of the TPC. Besides temperature fluctuations, a decrease in *E* with absolute latitude could also be explained by the metabolic cold adaptation hypothesis [52–55]. According to it, cold-adapted species should evolve lower thermal sensitivities (as well as higher *B*_0_ values; see Fig. 1) to maintain sufficient trait performance at very low temperatures. While our datasets do not possess the necessary resolution (especially at high latitudes; Figs. S11 and S13) for differentiating between these two alternative (and non-mutually exclusive) processes, this question remains to be addressed by future research.

Overall, a set of novel mechanistic explanations of TPC evolution emerge from our comparison of phylogenetic heritabilities of TPC parameters (Fig. 2). Contrary to *E* and *W*_op_, which have intermediate phylogenetic heritabilities, *T*_pk_ is almost perfectly phylogenetically heritable and evolves relatively gradually (i.e., without large jumps in parameter space; see Fig. S7 in the S1 Appendix). Thus, we expect TPCs to adapt to different thermal environments through both gradual changes in *T*_pk_ and discontinuous changes in *E*. Gradual changes in *T*_pk_ may be achieved through evolutionary shifts in the melting temperature of enzymes, i.e., the temperature at which 50% of the enzyme population is deactivated [56,57]. In contrast, changes in thermal sensitivity may be the outcome of i) evolution of enzymes with different heat capacities [57–59], ii) changes in the plasticity of cellular membranes [3,60], or even iii) restructuring of the underlying metabolic network [61].

Fundamental differences in the selection mechanisms underlying the evolution of *T*_pk_ and *E* may also explain the difference in evolutionary patterns between them. Specifically, both the mean environmental temperature (to which *T*_pk_ responds [7,35]) and the temperature fluctuations (to which *E* responds [3,21,24–26,35]) vary systematically from the equator to intermediate latitudes. We hypothesize that a species adapted to low temperatures is unlikely to adapt to a high-temperature environment rapidly enough (i.e., through a large increase in *T*_pk_) as it is pushed to its thermal tolerance limits [62,63]. In contrast, a species adapted to a fluctuating thermal environment (i.e., with a low *E* value) should be able to survive in more thermally stable conditions without much cost, becoming a thermal specialist (i.e., with a high *E* value) relatively rapidly, resulting in the observed jumps in trait space when mapped on the phylogeny (Figs. 3, 6, S9, and S10).

It is worth stressing, however, that not all types of thermal fluctuations are expected to impose selection for thermal generalists. In particular, thermal generalist variants of a given species are expected to be favoured when temperature fluctuations primarily occur across generations [24, 25, 64]. In contrast, moderate to strong thermal variation within generations would lead to selection for thermal specialists, even when inter-generational fluctuations are also present. For the microbial groups of the present study, an estimate of the minimum generation time can be calculated as the inverse of the *B*_pk_ of *r*_max_. Across our datasets of phytoplankton and prokaryotes, the minimum generation time ranges from a few minutes to 3.5 months, with phylogenetically-corrected medians of ≈ 40.5 hours for phytoplankton and ≈ 3.5 hours for prokaryotes (Fig. S14). Given this and because the magnitudes of annual and intra-annual (e.g., monthly) thermal fluctuations increase from the equator to intermediate latitudes [3,45,65], most microbes from intermediate latitudes are expected to generally experience substantial inter-generational thermal fluctuations and to a much lesser extent intra-generational fluctuations. This is indeed consistent with the observed weak decline in *E* at intermediate latitudes compared to the equator (Figs. 7 and S13). Nevertheless, latitude, trait identity, and species identity account for only 58% of the variance in *E*, indicating that adaptive shifts in *E* may also be driven by other factors such as biotic interactions [18, 66, 67]. A systematic identification of drivers of thermal sensitivity as well as the magnitude of their respective influence could be the focus of future studies.

For the thermal sensitivity of *r*_max_ in particular, the observed patterns of discontinuous evolution likely reflect the evolution of TPCs of underlying physiological traits on which it depends. For example, in populations of photosynthetic cells, shifts in the thermal sensitivity of any or all of photosynthesis rate, respiration rate, and carbon allocation efficiency can induce large changes in the *E* of *r*_max_ [30]. Indeed, we observed large adaptive shifts in thermal sensitivity even for fundamental physiological traits such as respiration rate (Fig. S10B,D), contrary to the MTE expectation of strong evolutionary conservatism [8–10]. This result is in agreement with a previous study that had identified significant adaptive variation in the thermal performance curve of the specific activity of Rubisco carboxylase [31]. It remains to be seen whether a similar lack of evolutionary conservation can be detected in key enzymes of non-photosynthetic organisms. Further research is clearly also needed on how the thermal sensitivities of different traits underlying fitness interact, and the extent to which these interactions can be modified through adaptation.

Besides biological-driven variation in thermal sensitivity, “artificial” variation may also be present, hindering the recognition of real patterns. For example, *E* estimates can be inaccurate if trait measurements in the rise of the TPC are limited, and span too narrow a range of temperatures [12]. To address this issue, we only kept *E* estimates if at least four trait measurements were available at the rise of each TPC. Further variation in thermal sensitivity can be introduced if trait values are measured instantaneously (without allowing sufficient time for acclimation) or under suboptimal conditions (e.g., under nutrient- or light-deficient conditions). Such treatments can lead to systematic biases in the shape of the resulting TPCs, which may strongly differ from TPCs obtained after adequate acclimation and under optimal growth conditions [27,68–71]. To avoid such biases, the datasets that we used only included TPCs that were experimentally determined after acclimation and under optimal conditions. On the other hand, maintenance of a given strain under a fixed set of experimental conditions for hundreds of generations could also lead to adaptive changes in TPC shape, due to the emergence of novel genetic mutations, as has been previously shown [26,27]. While the strains in our dataset were not grown over such long time periods, future studies could employ experimental evolution to measure the rate of thermal sensitivity evolution over much shorter time scales than the ones in our study.

Put together, all these results yield a compelling mechanistic explanation of how evolution shapes the distribution of *E*, and emphasize the need to consider the ecological and evolutionary underpinnings as well as implications of variation in *E*, as has been pointed out in a spate of recent studies [12,18,20,21,30]. In particular, our study helps explain the reason for the right skewness in the *E* distributions previously identified across practically all traits and taxonomic groups [12,18,21]. A clear explanation for this pattern was previously lacking, partly because MTE posits that *E* should be thermodynamically constrained and thus almost invariable across species [8–10]. Our study fills this gap in understanding by showing that the distribution of *E* is the outcome of frequent convergent evolution, driven by the adaptation of species from different clades to similar environmental conditions. In other words, as species encounter new environments through active or passive dispersal [72–74], they face selection for particular values of thermal sensitivity, which results in (often large) shifts in *E*. This process explains both the low variation in *E* among some species groups (Fig. 4) and the shape of its distribution. More precisely, the high degree of right skewness probably reflects the fact that most environments select for thermal generalists, with high *E* values being less frequently advantageous. Our findings have implications for ecophysiological models which may benefit from accounting for variation in thermal sensitivity among species or individuals. This could both yield an improved fit to empirical datasets [75] and provide a more realistic approximation of the processes being studied. Finally, the existence of adaptive variation in thermal sensitivity is likely to partly drive ecological patterns at higher scales (e.g., the response of an ecosystem to warming). How differences in thermal sensitivity among species influence ecosystem function is largely unaddressed [32,75] but highly important for accurately predicting the impacts of climate change on diverse ecosystems.

## Methods

### Phylogeny reconstruction and relative time calibration

We performed sequence alignment using MAFFT (v. 7.123b) [76] and its L-INS-i algorithm, and ran Noisy (v. 1.5.12) [77] with the default options to identify and remove phylogenetically uninformative homoplastic sites. For a more robust phylogenetic reconstruction, we used the results of previous phylogenetic studies by extracting the Open Tree of Life [78] topology for the species in our dataset using the rotl R package [79]. We manually examined the topology to eliminate any obvious errors. In total, 497 species were present in the tree, whereas many nodes were polytomic. To add missing species and resolve polytomies, we inferred 1,500 trees with RAxML (v. 8.2.9) [80] from our concatenated sequence alignment, using the Open Tree of Life topology as a backbone constraint and the General Time-Reversible model [81] with Γ-distributed rate variation among sites [82]. This model was fitted separately to each gene partition (i.e., one partition for the alignment of the small subunit rRNA gene sequences and one partition for the alignment of cbbL/rbcL gene sequences). Out of the 1,500 resulting tree topologies, we selected the tree with the highest log-likelihood and performed bootstrapping (using the extended majority-rule criterion) [83] to evaluate the statistical support for each node.

Finally, we calibrated the resulting RAxML tree to units of relative time by running DPPDiv [84] on the alignment of the small subunit rRNA gene sequences using the uncorrelated Γ-distributed rates model [85] (Fig. S1 in the S1 Appendix). For this, we used the alignment of small subunit rRNA gene sequences only, as DPPDiv can only be run on a single gene partition. We executed two DPPDiv runs for 9.5 million generations, sampling from the posterior distribution every 100 generations. After discarding the first 25% of samples as burn-in, we ensured that the two runs had converged on statistically indistinguishable posterior distributions by examining the effective sample size and the potential scale reduction factor [86,87] for all model parameters. More precisely, we verified that all parameters had an effective sample size above 200 and a potential scale reduction factor value below 1.1. To summarise the posterior distribution of calibrated trees into a single relative chronogram, we kept 4,750 trees per run (one tree every 1,500 generations) and calculated the median height for each node using the TreeAnnotator program [88].

### Sharpe-Schoolfield model fitting

To obtain estimates of the parameters of each experimentally determined TPC, we fitted the following four-parameter variant of the Sharpe-Schoolfield model (Fig. 1) [5, 35]:

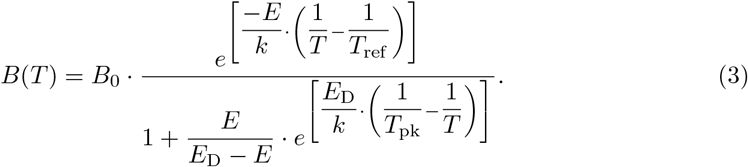

This model extends the Boltzmann-Arrhenius model (Eq. 1) to capture the decline in trait performance after the TPC reaches its peak (*T*_pk_). We followed the same approach for fitting the Sharpe-Schoolfield model as in reference [35]. Briefly, we set *T*_ref_ to 0°C, as for *B*_0_ to be biologically meaningful (see Fig. 1), it needs to be normalised at a temperature below the minimum *T*_pk_ in the study. Thus, a *T*_ref_ value of 0°C allowed us to include TPCs from species with low *T*_pk_ values in the analyses. Also, as certain specific TPC parameter combinations can mathematically lead to an overestimation of *B*_0_ compared to the true value, *B*(*T*_ref_) [89], we manually recalculated *B*(*T*_ref_) for each TPC after obtaining estimates of the four main parameters (*B*_0_, *E*, *T*_pk_, and *E*_D_). For simplicity, these recalculated *B*(*T*_ref_) values are referred to as *B*_0_ throughout the study. Finally, *B*_pk_ and *W*_op_ were calculated based on the estimates of the four main parameters.

After rejecting fits with an *R*^2^ below 0.5, there were i) 312 fits across 118 species from the phytoplankton *r*_max_ dataset, ii) 289 fits across 189 species from the prokaryote *r*_max_ dataset, iii) 87 fits across 38 species from the net photosynthesis rates dataset, and 34 fits across 18 species from the respiration rates dataset. Note that some species were represented by multiple fits due to the inclusion of experimentally-determined TPCs from different strains of the same species or from different geographical locations. To ensure that all TPC parameters were reliably estimated, we performed further filtering based on the following criteria: i) *B*_0_ and *E* estimates were rejected if fewer than four experimental data points were available below *T*_pk_. ii) Extremely high *E* estimates (i.e., above 4 eV) were rejected. iii) *W*_op_ values were retained if at least four data points were available below *T*_pk_ and two after it. iv) Two data points below and after the peak were required for accepting the estimates of *T*_pk_ and *B*_pk_. v) *E*_D_ estimates were kept if at least four data points were available at temperatures greater than *T*_pk_.

### Estimation of phylogenetic heritability for all TPC parameters

As in the previous subsection, the methodology that we used here was identical to that in reference [35]. In short, we specified a phylogenetic mixed-effects model for each of the two large TPC datasets with MCMCglmm. The models had a combined response with all TPC parameters transformed towards normality. The uncertainty for each estimate was obtained with the delta method [90] or via bootstrapping (for ln(*W*_op_)) and was incorporated into the model. Missing estimates in the response variables (i.e., when not all parameter estimates could be obtained for the same TPC) were modelled according to the “Missing At Random” approach [36,91]. Regarding fixed effects, a separate intercept was specified for each TPC parameter. Species identity was treated as a random effect on the intercepts and was corrected for phylogeny through the integration of the inverse of the phylogenetic variance-covariance matrix. We note that our approach is superior to the estimation of phylogenetic signal (Pagel’s *λ* [92]) separately for each TPC parameter, as the latter cannot account for covariance among TPC parameters, multiple measurements per species, or missing values. For each dataset, two Markov chain Monte Carlo chains were run for 200 million generations and estimates of the parameters of the model were sampled every 1,000 generations after the first 20 million generations were discarded as burn-in. Tests to ensure that the chains had converged and that the parameters were adequately sampled were done as previously described.

### Disparity-through-time analyses

We performed disparity-through-time analyses for ln(*E*) and ln(*W*_op_), using the rank envelope method [38] to generate a confidence envelope from 10,000 simulations of random evolution (Brownian motion). As it is not straightforward to incorporate multiple measurements per species with this method, we selected the ln(*E*) or ln(*W*_op_) estimate of the Sharpe-Schoolfield model fit with the highest *R*^2^ value per species.

### Free, stable, and Lévy model fitting

We fitted the free, stable, and Lévy models of trait evolution to estimates of ln(*E*) and ln(*W*_op_), using the motmot.2.0 R package (v. 1.1.2) [93,94], the stabletraits software [42], and the levolution software [43] respectively. To obtain each fit of the stable model, we executed four independent Markov chain Monte Carlo chains for 30 million generations, recording posterior parameter samples every 100 generations. Samples from the first 7.5 million generations were excluded, whereas the remaining samples were examined to ensure that convergence had been achieved. For fitting the Lévy model, we used the peak-finder algorithm to estimate the value of the model’s *α* parameter. More precisely, we set the starting value of *α* to 10^0.5^, the step size to 0.5, and the number of optimizations to 5, as suggested in levolution’s documentation. We also changed the maximum number of iterations (option “`-maxIterations`”) to 2,000 so that the algorithm could sufficiently converge in all cases.

### Investigation of a putative relationship between latitude and ln(E) and ln(W_op_)

We examined the relationship of thermal sensitivity with latitude by fitting regression models with MCMCglmm to all four TPC datasets combined. The response variable was ln(*E*) or ln(*W*_op_), whereas possible predictor variables were i) latitude (in radian units and using a cosine transformation, as absolute latitude in degree units, or split in three bins of low, intermediate, and high absolute latitude; subsections S4.2 and S4.3 in the S1 Appendix), ii) the trait from which thermal sensitivity estimates were obtained, and iii) the interaction between latitude and trait identity. To properly incorporate multiple measurements from the same species (where available), we treated species identity as a random effect on the intercept. We fitted both phylogenetic and non-phylogenetic variants of all candidate models. Two chains per model were run for five million generations each, with samples from the posterior being captured every thousand generations. We verified that each pair of chains had sufficiently converged, after discarding samples from the first 500,000 generations. To identify the most appropriate model, we first rejected models that had a non-intercept coefficient with a 95% Highest Posterior Density (HPD) interval that included zero. We then selected the model with the lowest mean DIC value. To report the proportions of variance explained by the fixed effects (Var_fixed_), by the random effect (Var_random_), or left unexplained (Var_resid_), we calculated the marginal and conditional coefficients of determination [95]:

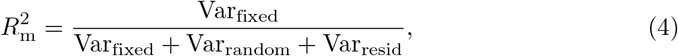

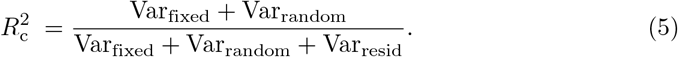

### Mantel test between phylogenetic and latitudinal distance matrices

We used the R package ade4 (v. 1.7-13) [96] to infer the correlation of phylogenetic distance with latitudinal distance across phytoplankton and prokaryotes using the Mantel test. To generate the *P*-values, we set the number of permutations to 9,999.

## Acknowledgments

We thank Iain Colin Prentice for providing comments on an early version of the manuscript. We are also grateful to James Rosindell, Jonathan Lloyd, Guy Woodward, and Andrew G. Hirst for valuable discussions, and to the CIPRES Science Gateway [97] for access to computational resources. Species silhouettes were obtained from phylopic.org and are used under the Public Domain license. The images of species in Fig. 1 were graciously provided by Eric Erbe, Christopher Pooley, and the Rocky Mountain National Park, also under the Public Domain license. DGK was supported by a Natural Environment Research Council (NERC) Doctoral Training Partnership (DTP) scholarship (NE/L002515/1). TPS was supported by a Biotechnology and Biological Sciences Research Council (BBSRC) DTP scholarship (BB/J014575/1). SP was supported by NERC grants NE/M004740/1 and NE/M020843/1.

