## Supplementary material for "Adaptive evolution shapes the present-day distribution of the thermal sensitivity of population growth rate": S1 Appendix

**S5 Minimum generation times of microbes** **13**

**S6 List of nucleotide sequences used for phylogeny reconstruction** **13**

**1** Science and Solutions for a Changing Planet DTP, Imperial College London, London, United Kingdom.
**2** Department of Life Sciences, Imperial College London, Silwood Park, Ascot, Berkshire, United Kingdom.

### S1 Phylogeny reconstruction

The final tree produced by RAxML [1] and calibrated to relative time with DPPDiv [2] is shown in Fig. S1.

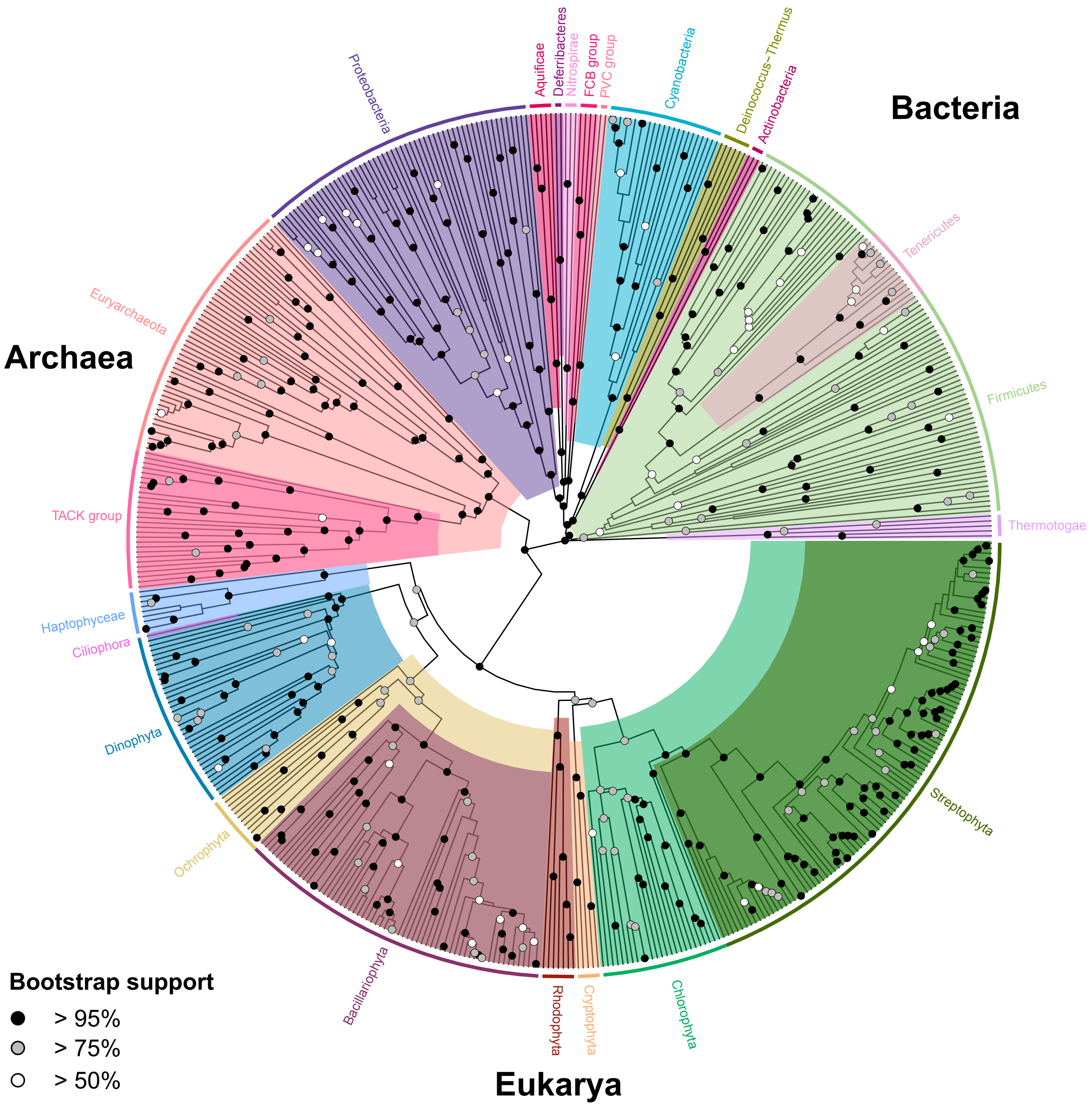

**Figure S1. The phylogeny generated in this study from which subtrees were extracted for comparative analyses.** Colours indicate different phyla, whereas circles show the statistical support for each node, conditional to the topological constraints of the Open Tree of Life [3].

#### S2 Dataset of thermal sensitivity estimates

The distributions of $E$ and $W_{op}$ values across the four datasets for species included in the phylogeny are shown in Fig. S2. Fig. S3 shows the distributions of thermal sensitivity estimates of $r_{max}$ across the six largest phyla of this study.

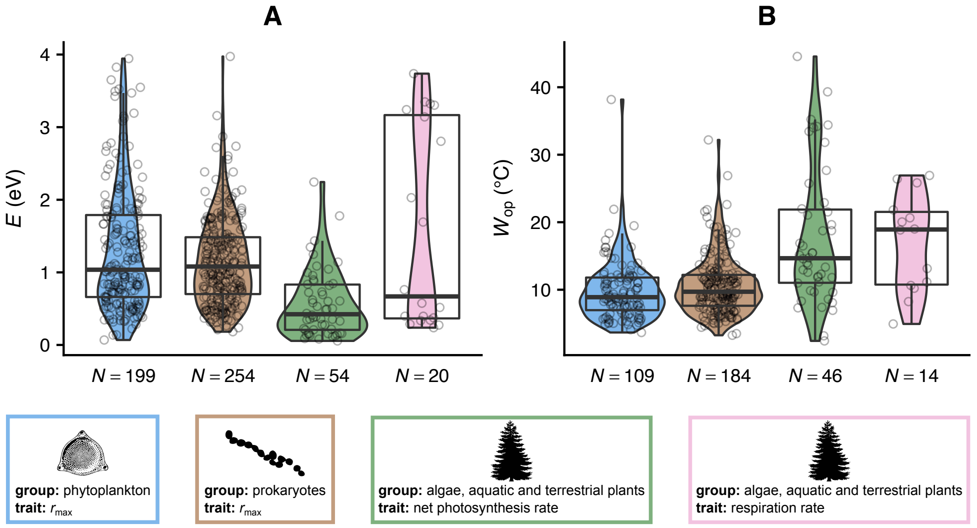

**Figure S2. All estimates of thermal sensitivity that were part of this study.** Multiple estimates from the same species are not averaged but are included as separate data points. As experimentalists rarely measure trait performance across the entire TPC, for many TPCs it was not possible to robustly estimate both $E$ and $W_{op}$. For this reason, $E$ and $W_{op}$ do not have the same sample size for each dataset. Overall, the distributions of thermal sensitivity parameters are not approximately Gaussian but asymmetric and not generally inflated at their boundaries. This indicates that the variation in thermal sensitivity is real and not purely due to measurement error.

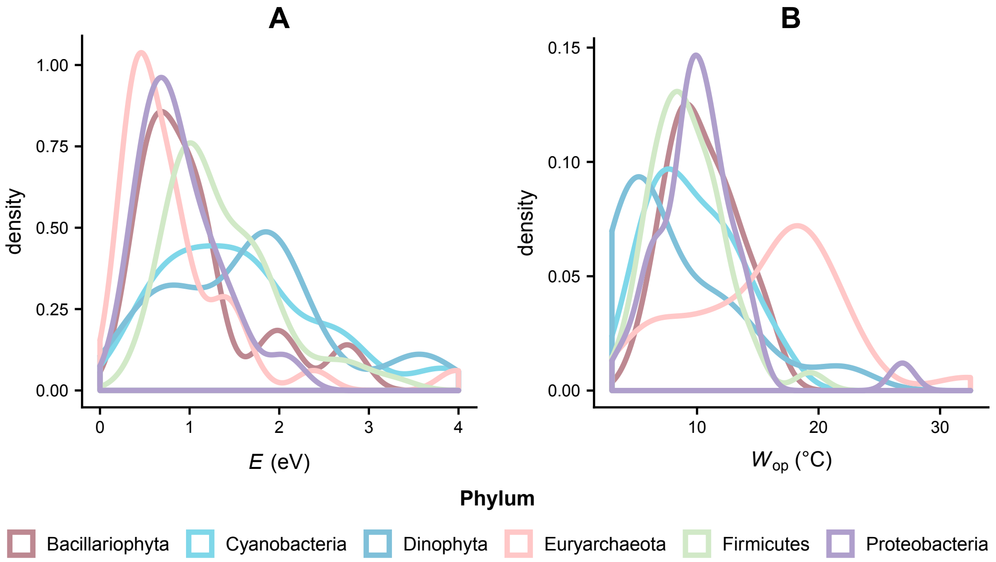

**Figure S3. The $E$ and $W_{op}$ distributions of the largest phyla in the study exhibit considerable overlap.** Here, each species is represented by a single thermal sensitivity estimate. The increasing convergence of thermal sensitivity distributions even among evolutionarily remote phyla (e.g., between Cyanobacteria and Dinophyta or between Bacillariophyta and Proteobacteria) explains the intermediate phylogenetic heritability of thermal sensitivity. Moreover, it suggests that different values of $E$ or $W_{op}$ correspond to distinct thermal strategies which species can evolve largely regardless of their evolutionary background.

### S3 Phylogenetic comparative analyses

#### S3.1 Analyses of the datasets of $r_{max}$ TPCs

For the estimation of phylogenetic heritability for each TPC parameter, we inferred the variance/covariance matrix of TPC parameters, corrected for phylogeny. This allowed us to also extract the phenotypic correlation ($r_{phe}$) between $E$ and $W_{op}$ and, thus, to understand the relationship between the two thermal sensitivity measures (Fig. S4). Furthermore, the phenotypic correlation was broken down to its phylogenetically heritable component ($r_{her}$) and its residual component ($r_{res}$). The latter should be driven mostly by environmental effects.

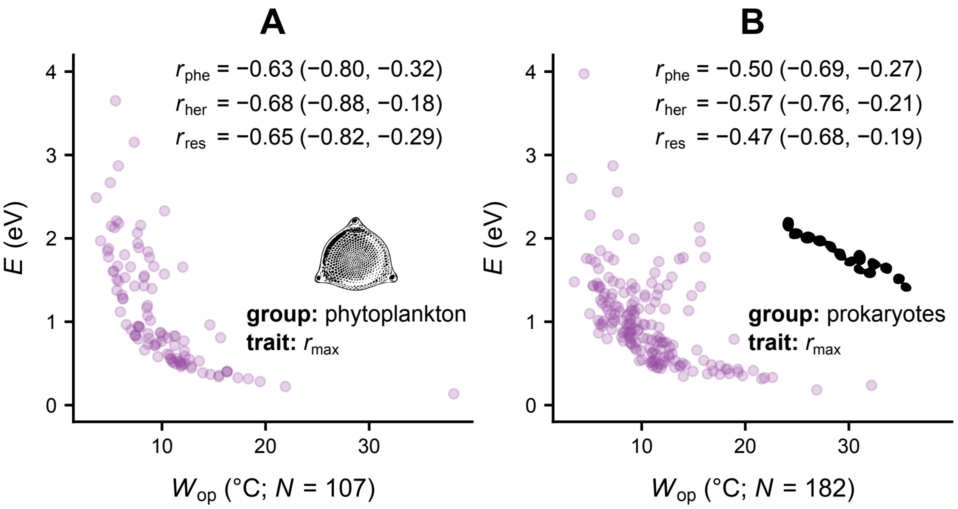

**Figure S4. Correlations between $E$ and $W_{op}$ among phytoplankton and prokaryotes.** Despite both being measures of thermal sensitivity, $E$ and $W_{op}$ are not perfectly correlated. The correlation coefficients shown are posterior distribution means, with values in parentheses indicating the 95% Highest Posterior Density interval. All correlation estimates were obtained for $\ln(E)$ and $\ln(W_{op})$, whereas here the parameters are shown untransformed.

To examine how the evolutionary rate of thermal sensitivity varies across the phylogeny, we fitted the stable model of trait evolution [4] (Fig. 5 in the main text) but also the free model [5] (Fig. S5) and the Lévy model [6] (Fig. S6).

Finally, to better understand how species explore the parameter space of $E$, $W_{op}$, and $T_{pk}$ (whose phylogenetic heritability is $\approx 1$; see Fig. 2), we combined our two $r_{max}$ datasets and divided the distributions of $E$, $W_{op}$, and $T_{pk}$ into four discrete states (Fig. S7). Boundaries for these states were selected using the Jenks natural breaks clustering algorithm [7], as implemented in the BAMMtools R package (v. 2.1.6) [8]. To estimate the transition rates among states, we fitted the "all-rates-different" variant of the M$k$ model [9] with the `fitMk` function of the phytools R package (v. 0.6-60) [10].

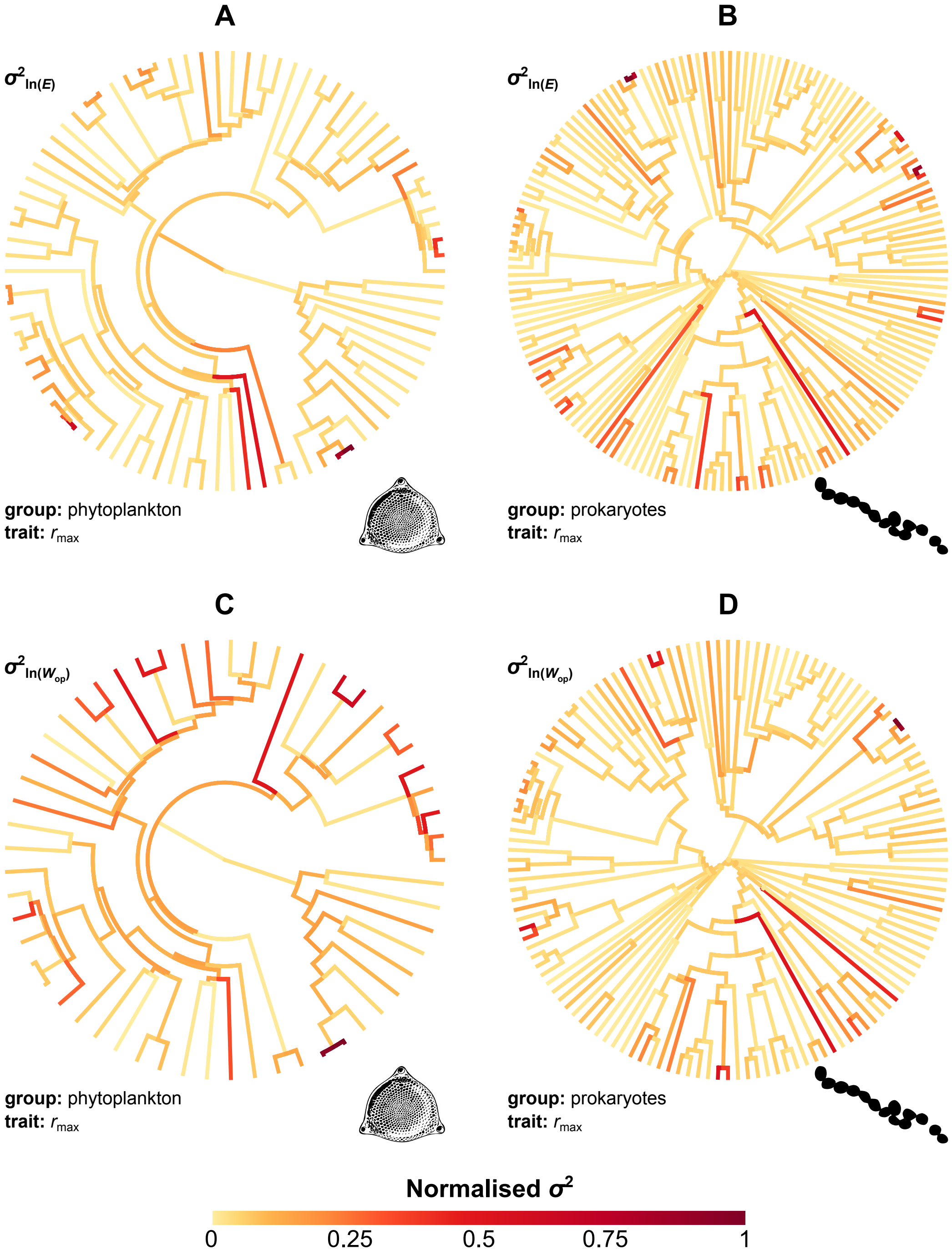

**Figure S5. Variation in the evolutionary rate across the phylogeny, as inferred with the free model.** The results are qualitatively similar to those obtained with the stable model. The highest evolutionary rates (dark red and brown) generally appear in late-branching lineages across the phylogeny and are not clustered in specific clades.

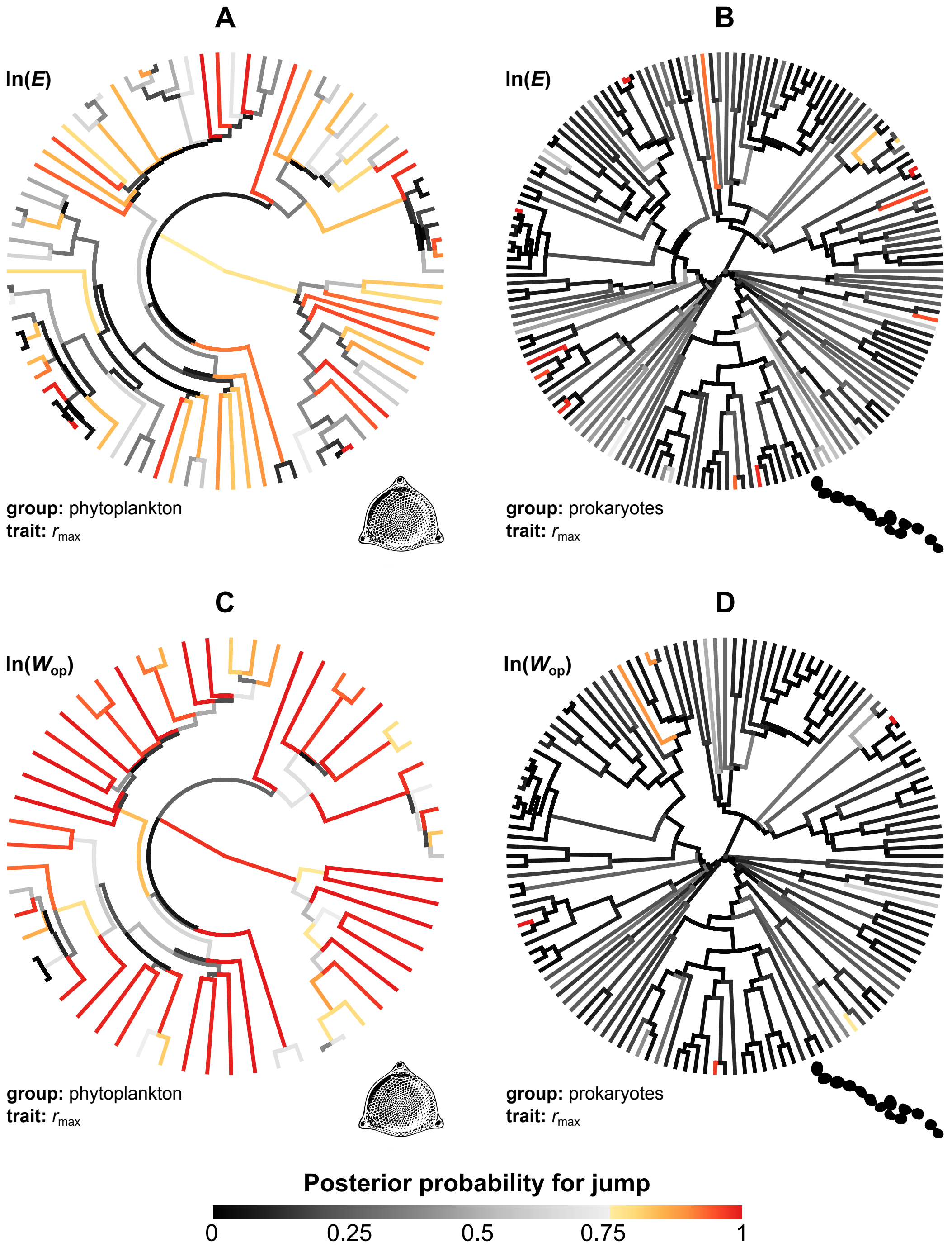

**Figure S6. Evolutionary jumps in thermal sensitivity across the phylogeny, as inferred with the Lévy model.** The results are again qualitatively similar to those obtained with the stable model. Branches with a high posterior probability for the occurrence of a jump (shown in yellow to dark red) are distributed across the entire phylogeny and are not limited to specific clades. Note that in all cases, the Lévy model had a much lower AIC (between 20 and 147 units difference) than the constant-rate Brownian motion model (the null expectation).

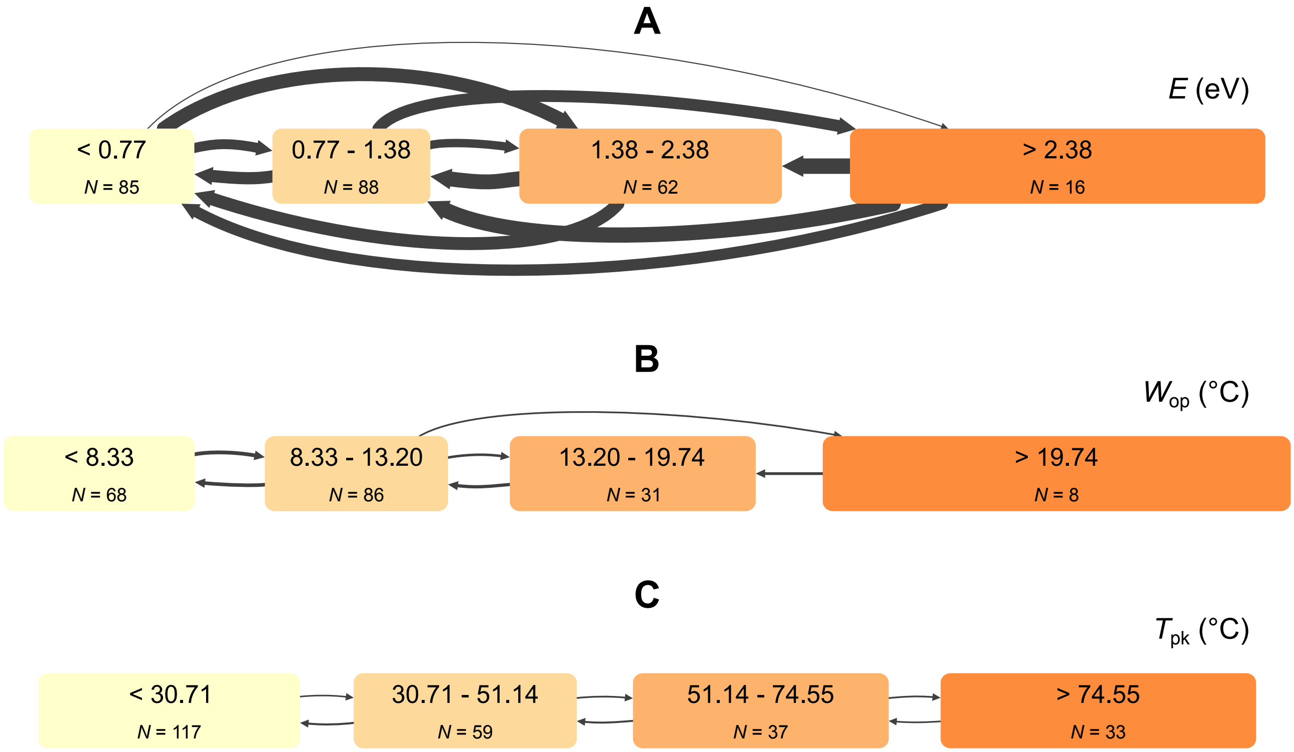

**Figure S7. Transitions in the discretized parameter space of $E$, $W_{\mathbf{op}}$, and $T_{\mathbf{pk}}$.** The width of the edges represents the natural logarithm of the transition rate between states. Transitions between non-neighbouring states are very common for $E$ (which captures the rise of the TPC), rare for $W_{\mathrm{op}}$ (which captures both the rise and the peak of the TPC), and never observed for $T_{\mathrm{pk}}$ (which captures the peak of the TPC). It is worth pointing out that $T_{\mathrm{pk}}$ also exhibits the lowest transition rates between neighbouring states among the three TPC parameters. These results are consistent with the phylogenetic heritability estimates shown in Fig. 2 in the main text.

#### S3.2 Analyses of the dataset of phytoplankton TPCs after excluding Cyanobacteria

Removing Cyanobacteria from the phytoplankton dataset led to qualitatively identical results in our phylogenetic analyses (Figs. S8 and S9).

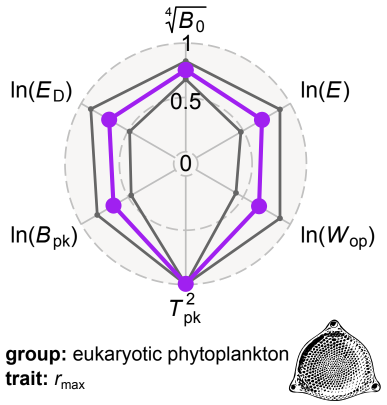

**Figure S8. Phylogenetic heritabilities of TPC parameters of eukaryotic phytoplankton.** The main differences between these results and those using the entire phytoplankton dataset (Fig. 2A in the main text) were that, here, $\ln(E)$ and $\ln(B_{pk})$ have slightly higher/lower phylogenetic heritabilities respectively. The former is expected as the thermal sensitivity distribution of Cyanobacteria is very similar to that of Dinophyta (Fig. 4 in the main text), despite the long evolutionary distance between them. Therefore, the exclusion of Cyanobacteria would necessarily increase the phylogenetic heritability of thermal sensitivity. Similarly, $\ln(B_{pk})$ in prokaryotes is more phylogenetically heritable than in phytoplankton (Fig. 2 in the main text), explaining the further decrease in its phylogenetic heritability when Cyanobacteria are excluded.

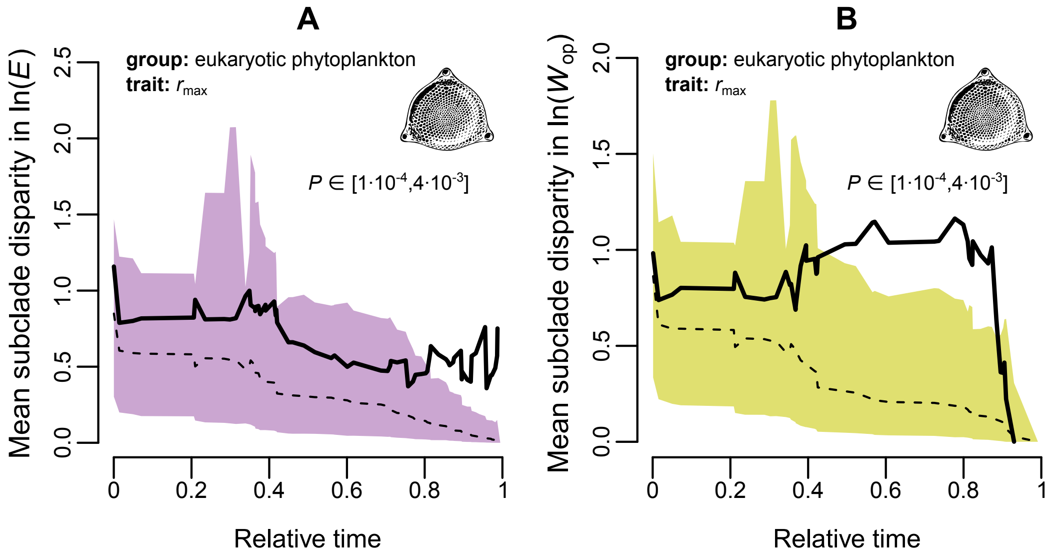

**Figure S9. The mean subclade disparity in thermal sensitivity tends to increase with time across phytoplankton, even after excluding Cyanobacteria.** The pattern is slightly weaker (albeit still present) due to the lower sample size.

#### S3.3 Analyses of the net photosynthesis rate and respiration rate TPC datasets

The visualization of the evolution of thermal sensitivity of net photosynthesis rate and respiration rate revealed similar patterns to those of the thermal sensitivity of $r_{\text{max}}$ (Fig. 6 in the main text). Thermal sensitivity values do not evolve gradually and tightly around a central value ($\widehat{\theta}$), but explore large parts of the parameter space due to bursts of rapid evolution.

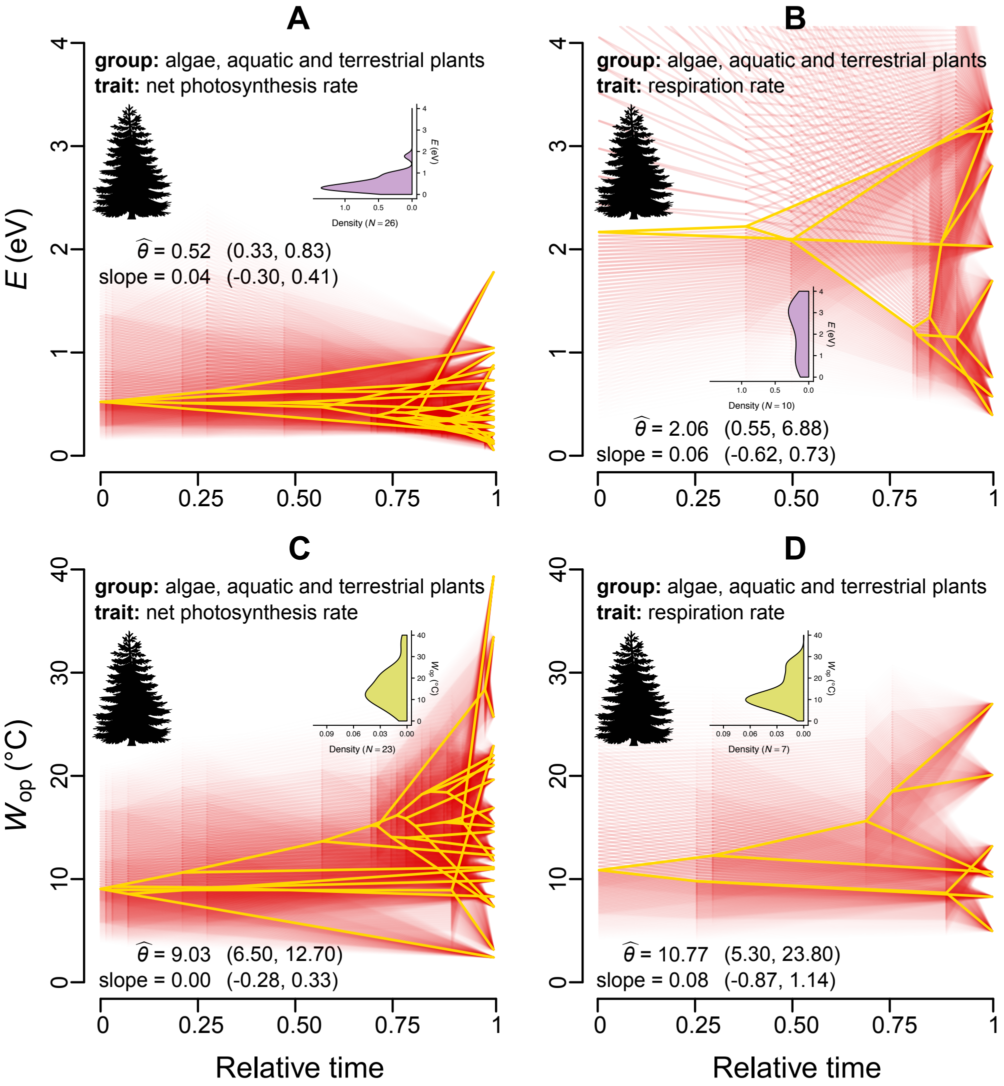

**Figure S10. Evolution of the thermal sensitivities of net photosynthesis rate and respiration rate through time.** The inset figures show the density distributions of $E$ and $W_{\text{op}}$ values of extant species in the dataset.

### S4 Investigation of latitudinal associations for measures of thermal sensitivity

#### S4.1 Latitudinal coverage of the dataset

The latitudes of species/strains from which we obtained estimates of thermal sensitivity ($E$ and $W_{op}$) are shown in Fig. S11.

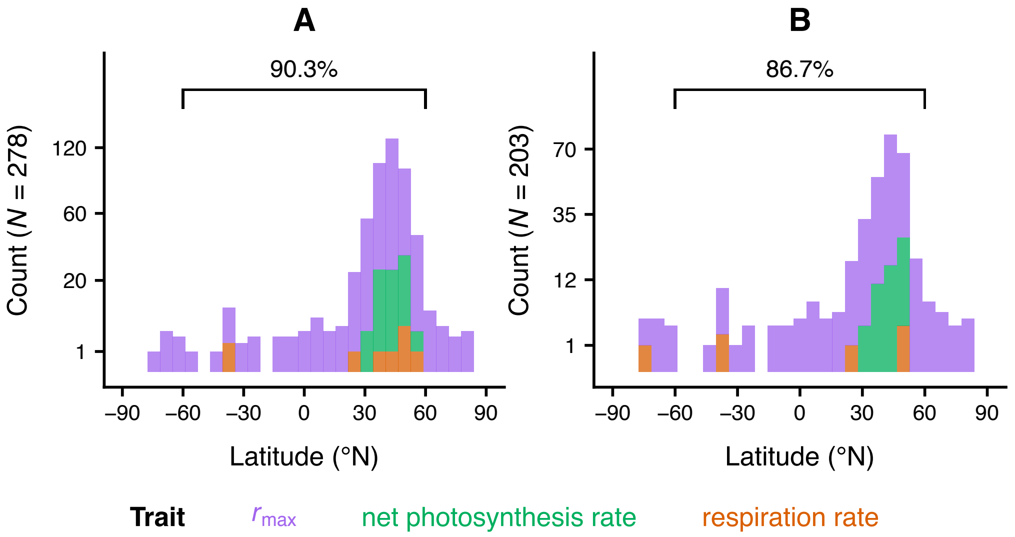

**Figure S11. Latitudinal distribution of $E$ (A) and $W_{op}$ (B) estimates in this study.** Most estimates are from species/strains found at low and intermediate latitudes. The percentage at the top of each panel indicates the fraction of estimates from latitudes between -60°N and 60°N. Note that values along the vertical axes do not increase linearly.

#### S4.2 Fitted models using latitude as a continuous predictor

We rejected models that had one or more non-intercept coefficients with a 95% HPD interval that included zero. We then used DIC to identify the most appropriate model among those remaining.

| Model | Phylogenetic correction | Mean DIC |
|---|---|---|
| $\ln(E) \sim$ Intercept + cos(Latitude) + Trait identity | ✓ | 535.2840 |
| $\ln(E) \sim$ Intercept + \|Latitude\| + Trait identity | ✓ | 536.0311 |
| $\ln(E) \sim$ Intercept + Trait identity | ✓ | 538.7041 |
| $\ln(E) \sim$ Intercept | ✓ | 569.9043 |
| $\ln(E) \sim$ Intercept + cos(Latitude) + Trait identity | ✗ | 489.5730 |
| $\ln(E) \sim$ Intercept + \|Latitude\| + Trait identity | ✗ | 488.8540 |
| $\ln(E) \sim$ Intercept + cos(Latitude) | ✗ | 511.4035 |
| $\ln(E) \sim$ Intercept + \|Latitude\| | ✗ | 510.1707 |
| $\ln(E) \sim$ Intercept + Trait identity | ✗ | 492.4570 |
| $\ln(E) \sim$ Intercept | ✗ | 514.0522 |

**Table S1. Candidate models with $\ln(E)$ as the response variable.**

| Model | Phylogenetic correction | Mean DIC |
|---|---|---|
| $\ln(W_{op}) \sim$ Intercept | ✓ | 148.6599 |
| $\ln(W_{op}) \sim$ Intercept + Trait identity | ✗ | 144.8427 |
| $\ln(W_{op}) \sim$ Intercept | ✗ | 147.4493 |

**Table S2. Candidate models with $\ln(W_{op})$ as the response variable.**

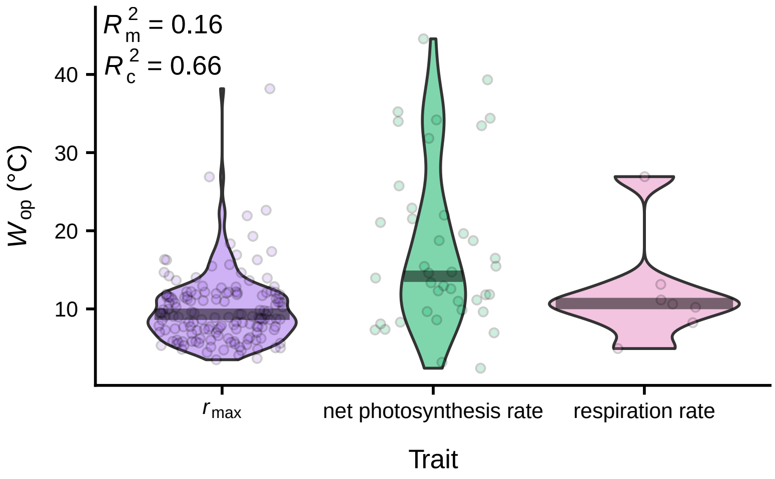

**Figure S12. The most appropriate model for $\ln(W_{op})$ does not include latitude as a predictor variable.** Trait identity on its own accounts for 16% of variance, whereas also incorporating species identity (as a random effect on the intercept) results in 66% of the variance being explained. Grey horizontal lines denote the inferred intercept for each trait.

#### S4.3 Fitted models using binned latitude as predictor

| Model | Phylogenetic correction | Mean DIC |
|---|---|---|
| $\ln(E) \sim$ Intercept + $\lvert\text{Latitude}\rvert_{0-30}$ + $\lvert\text{Latitude}\rvert_{>60}$ + Trait identity | ✓ | 535.8695 |
| $\ln(E) \sim$ Intercept + $\lvert\text{Latitude}\rvert_{0-30}$ + $\lvert\text{Latitude}\rvert_{>60}$ | ✓ | 565.1168 |
| $\ln(E) \sim$ Intercept + Trait identity | ✓ | 538.7041 |
| $\ln(E) \sim$ Intercept | ✓ | 569.9043 |
| $\ln(E) \sim$ Intercept + $\lvert\text{Latitude}\rvert_{0-30}$ + $\lvert\text{Latitude}\rvert_{>60}$ + Trait identity | ✗ | 485.0223 |
| $\ln(E) \sim$ Intercept + $\lvert\text{Latitude}\rvert_{0-30}$ + $\lvert\text{Latitude}\rvert_{>60}$ | ✗ | 504.8906 |
| $\ln(E) \sim$ Intercept + Trait identity | ✗ | 492.4570 |
| $\ln(E) \sim$ Intercept | ✗ | 514.0522 |

**Table S3. Candidate models for the effects of binned absolute latitude on $\ln(E)$.**

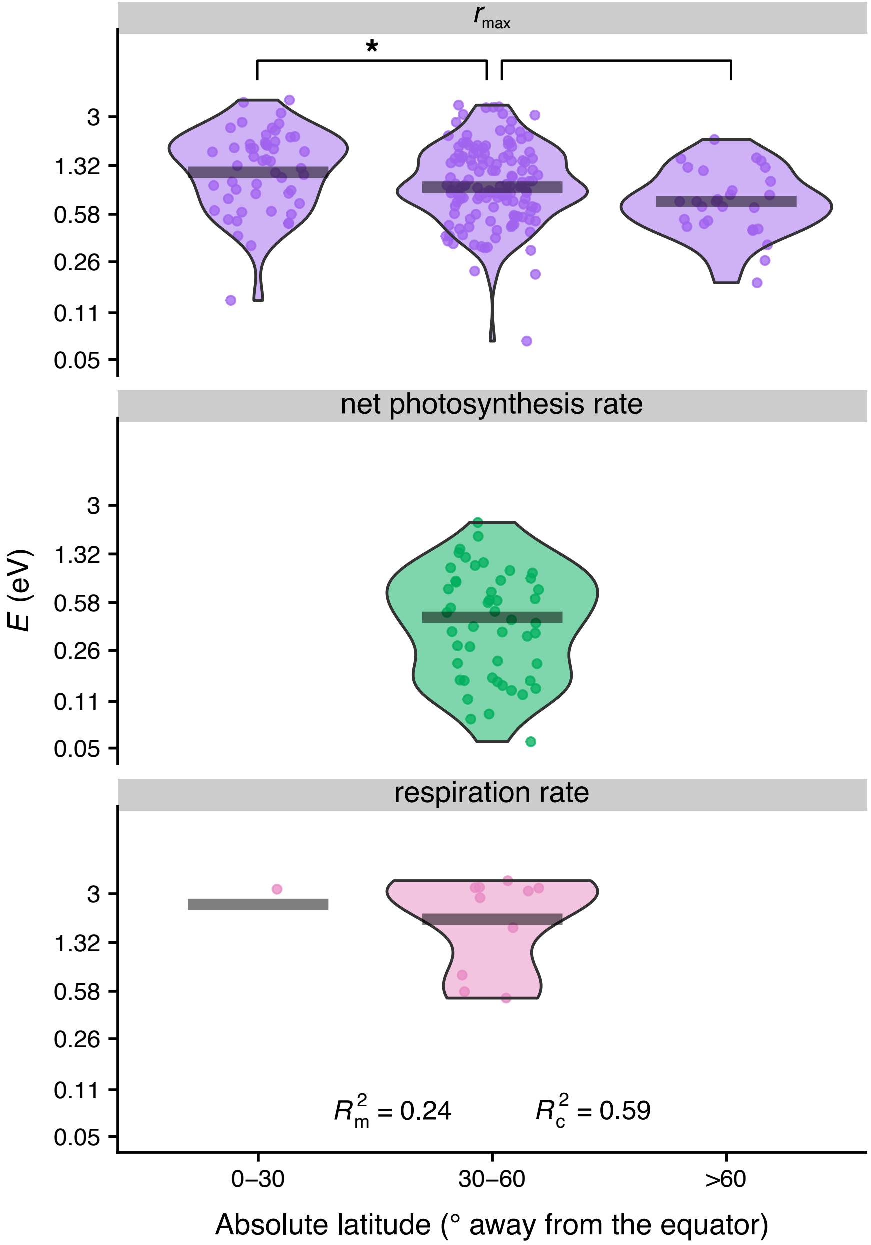

**Figure S13. Distribution of $E$ estimates across low, intermediate, and high latitudes.** Species/strains found in low latitudes tend to have slightly higher $E$ values than those in intermediate latitudes. In contrast, the $E$ distributions of intermediate and high latitudes were statistically indistinguishable.

### S5 Minimum generation times of microbes

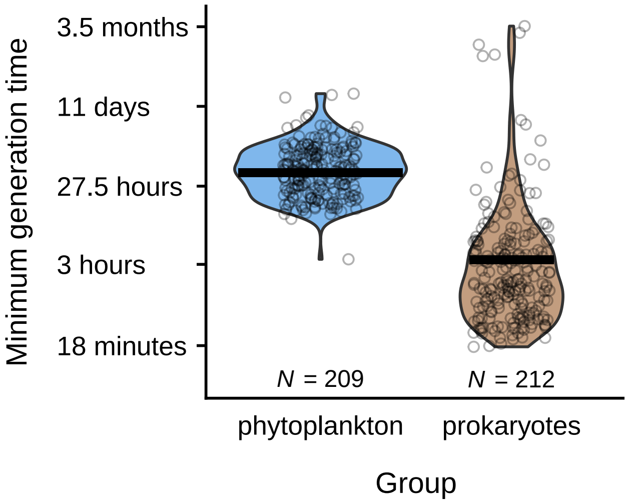

**Figure S14. Distribution of minimum generation time estimates for phytoplankton and prokaryotes.** Data points were obtained by taking the inverse of all $B_{pk}$ estimates. Horizontal bars represent the phylogenetically-corrected median values, i.e., the inverse of the intercept of $B_{pk}$ from the multi-response regression models that we fitted with MCMCglmm (see the "Estimation of phylogenetic heritability for all TPC parameters" subsection of the Methods in the main text).

### S6 List of nucleotide sequences used for phylogeny reconstruction

**Table S4.** Species names and Accession IDs of small subunit rRNA gene sequences that were used in this study.

| **Species** | **Accession ID** |
|---|---|
| *Abies alba* | GenBank: DQ371809.1 |
| *Abutilon theophrasti* | GenBank: DQ287985.1 |
| *Acer rubrum* | GenBank: U42494.1 |
| *Acetobacterium bakii* | NCBI Reference Sequence: NR_026329.1 |
| *Acetobacterium carbinolicum* | NCBI Reference Sequence: NR_026325.1 |
| *Acetobacterium fimetarium* | NCBI Reference Sequence: NR_026328.1 |
| *Acetobacterium paludosum* | NCBI Reference Sequence: NR_026327.1 |
| *Acetobacterium tundrae* | NCBI Reference Sequence: NR_028934.1 |
| *Acetogenium kivui* | NCBI Reference Sequence: NR_044617.1 |
| *Acidianus brierleyi* | NCBI Reference Sequence: NR_043409.1 |
| *Acidianus infernus* | NCBI Reference Sequence: NR_043431.1 |
| *Acidianus manzaensis* | GenBank: EU563854.1 |
| *Acidibacter ferrireducens* | NCBI Reference Sequence: NR_126260.1 |
| *Acidicaldus organivorus* | NCBI Reference Sequence: NR_042752.1 |
| *Acidilobus aceticus* | NCBI Reference Sequence: NR_041774.1 |
| *Acidilobus sulfurireducens* | NCBI Reference Sequence: NR_115940.1 |
| *Acidimicrobium ferrooxidans* | NCBI Reference Sequence: NR_074390.1 |
| *Acidithiobacillus caldus* | GenBank: KJ944319.1 |
| *Acidithiobacillus ferrivorans* | GenBank: KJ679874.1 |
| *Acidithiobacillus ferrooxidans* | GenBank: DQ062118.1 |
| *Acidithiobacillus thiooxidans* | GenBank: DQ834372.1 |
| *Acidocella aromatica* | GenBank: AF253413.1 |
| *Aeribacillus pallidus* | NCBI Reference Sequence: NR_026515.1 |

Table S4 – *Continued from previous page*

| Species | Accession ID |
|---|---|
| *Aeromonas hydrophila* | GenBank: M59148.1 |
| *Aeropyrum pernix* | NCBI Reference Sequence: NR_043417.1 |
| *Aldrovanda vesiculosa* | GenBank: AY096114.1 |
| *Alexandrium catenella* | GenBank: AJ535392.1 |
| *Alexandrium fundyense* | GenBank: KF908796.1 |
| *Alexandrium minutum* | GenBank: U27499.1 |
| *Alexandrium monilatum* | GenBank: AY883005.1 |
| *Alexandrium ostenfeldii* | GenBank: U27500.1 |
| *Alexandrium tamarense* | GenBank: AJ415510.1 |
| *Alkaliphilus transvaalensis* | GenBank: AJ630291.1 |
| *Amphidinium klebsii* | GenBank: EU046335.1 |
| *Amphiprora paludosa* | GenBank: AY485468.1 |
| *Anabaena bergii* | GenBank: AF160256.1 |
| *Anabaena macrospora* | GenBank: AJ293115.1 |
| *Anabaena spiroides* | GenBank: AB271212.1 |
| *Anabaena ucrainica* | GenBank: AB551452.1 |
| *Anabaena variabilis* | GenBank: AB016520.1 |
| *Ankistrodesmus falcatus var. tumidus* | GenBank: JQ315498.1 |
| *Antirrhinum majus* | GenBank: AJ236047.1 |
| *Aphanizomenon flosaquae* | GenBank: HE975013.1 |
| *Aphanizomenon gracile* | GenBank: AJ293127.1 |
| *Aphanizomenon ovalisporum* | GenBank: FM177484.1 |
| *Aplectrum hyemale* | GenBank: U59937.1 |
| *Arabidopsis thaliana* | NCBI Reference Sequence: NR_141642.1 |
| *Arbutus unedo* | GenBank: AF206853.1 |
| *Archaeoglobus veneficus* | NCBI Reference Sequence: NR_102885.1 |
| *Aristotelia serrata* | GenBank: GU476422.1 |
| *Asterionella formosa* | GenBank: AM712617.1 |
| *Asterionellopsis glacialis* | GenBank: X77701.1 |
| *Aulacoseira baicalensis* | GenBank: AY121821.1 |
| *Aulacoseira granulata* | GenBank: AB430586.1 |
| *Bacillus acidocaldarius* | GenBank: X60742.1 |
| *Bacillus caldotenax* | GenBank: AY608937.1 |
| *Bacillus cereus* | GenBank: KU198623.1 |
| *Bacillus infernus* | NCBI Reference Sequence: NR_027227.1 |
| *Bacillus megaterium* | GenBank: HM371417.1 |
| *Bacillus subtilis* | GenBank: AY728013.1 |
| *Beta vulgaris* | GenBank: FJ669720.1 |
| *Betula papyrifera* | GenBank: L00971.1 |
| *Betula pendula* | GenBank: GU476453.1 |
| *Brassica oleracea* | GenBank: KT225359.1 |
| *Brassica rapa* | GenBank: LC009534.1 |
| *Brochothrix thermosphacta* | GenBank: M58798.1 |
| *Bryum argenteum* | GenBank: U18529.1 |
| *Caldicellulosiruptor obsidiansis* | NCBI Reference Sequence: NR_117295.1 |
| *Caldisphaera draconis* | NCBI Reference Sequence: NR_115941.1 |
| *Caldivirga maquilingensis* | NCBI Reference Sequence: NG_042069.1 |
| *Caloramator indicus* | NCBI Reference Sequence: NR_026134.1 |
| *Caloranaerobacter azorensis* | NCBI Reference Sequence: NR_028919.1 |
| *Candidatus Brocadia sinica* | GenBank: KT023578.1 |
| *Capsella bursa-pastoris* | GenBank: KT459181.1 |
| *Capsicum annuum* | GenBank: EF564281.1 |

Table S4 – *Continued from previous page*

| Species | Accession ID |
|---|---|
| *Carya glabra* | GenBank: AF206880.1 |
| *Caulerpa serrulata* | GenBank: JQ745683.1 |
| *Ceratium furca* | GenBank: AJ276699.1 |
| *Ceratium fusus* | GenBank: AF022153.1 |
| *Ceratophyllum demersum* | GenBank: U42517.1 |
| *Chaetoceros debilis* | GenBank: AB847419.1 |
| *Chamaebatiaria millefolium* | GenBank: DQ886366.1 |
| *Chamerion angustifolium* | GenBank: AH001636.2 |
| *Chatonella marina* | GenBank: AB217627.1 |
| *Chenopodium album* | GenBank: HQ827790.1 |
| *Chlamydomonas reinhardtii* | GenBank: KF864473.1 |
| *Chlamydomonas subcaudata* | GenBank: AJ781310.1 |
| *Chlorella ellipsoidea* | GenBank: X63520.1 |
| *Chlorella pyrenoidosa* | GenBank: AB240151.1 |
| *Chlorella saccharophila* | GenBank: AB183577.1 |
| *Chlorella sorokiniana* | GenBank: EU402596.1 |
| *Chlorella vulgaris* | GenBank: HQ702325.1 |
| *Chlorobium tepidum* | NCBI Reference Sequence: NR_044685.2 |
| *Chondrus crispus* | GenBank: DQ317002.1 |
| *Chroomonas salina* | GenBank: GU983864.1 |
| *Chrysanthemum morifolium* | GenBank: KJ870235.1 |
| *Cicer arietinum* | GenBank: AJ011011.4 |
| *Citrus aurantium* | GenBank: U38312.1 |
| *Citrus limon* | GenBank: KJ740202.1 |
| *Cladophora glomerata* | GenBank: AB665579.1 |
| *Closterium acerosum* | GenBank: AF352230.1 |
| *Clostridium autoethanogenum* | NCBI Reference Sequence: NR_119283.1 |
| *Clostridium fervidus* | GenBank: L09187.1 |
| *Clostridium paradoxum* | NCBI Reference Sequence: NR_119327.1 |
| *Clostridium perfringens* | GenBank: LC037206.1 |
| *Clostridium thermoalcaliphilum* | GenBank: FR749953.1 |
| *Clostridium thermohydrosulfuricum* | NCBI Reference Sequence: NR_044618.1 |
| *Clostridium thermosuccinogenes* | GenBank: Y18180.1 |
| *Clostridium thermosulfurogenes* | GenBank: HG324062.2 |
| *Coccolithus pelagicus* | GenBank: AJ246261.1 |
| *Cochlodinium polykrikoides* | GenBank: EU418971.1 |
| *Coelastrum microporum* | GenBank: JQ315527.1 |
| *Colwellia demingiae* | NCBI Reference Sequence: NR_118860.1 |
| *Colwellia hornerae* | NCBI Reference Sequence: NR_118861.1 |
| *Colwellia psychrerythraea* | NCBI Reference Sequence: NR_037047.1 |
| *Colwellia psychrotropica* | NCBI Reference Sequence: NR_026055.1 |
| *Coolia monotis* | GenBank: EF492487.1 |
| *Coscinodiscus concinnus* | GenBank: HQ912681.1 |
| *Coscinodiscus granii* | GenBank: AY485495.1 |
| *Coscinodiscus jonesianus* | GenBank: KJ577852.1 |
| *Cosmarium biretum* | GenBank: AM920339.1 |
| *Cosmarium botrytis* | GenBank: AM920378.1 |
| *Cosmarium crenatum* | GenBank: AM920370.1 |
| *Cosmarium meneghinii* | GenBank: AM920366.1 |
| *Cosmarium punctulatum* | GenBank: AM920373.1 |
| *Cosmarium subprotumidum* | GenBank: AM920375.1 |
| *Cryptomonas erosa* | GenBank: AM396361.1 |

Table S4 – *Continued from previous page*

| Species | Accession ID |
|---|---|
| *Cryptomonas marssonii* | GenBank: EU163586.1 |
| *Cryptomonas ovata* | GenBank: KC928318.1 |
| *Cucumis sativus* | GenBank: AF206894.1 |
| *Cyclotella cryptica* | GenBank: AY485499.1 |
| *Cyclotella meneghiniana* | GenBank: HM805030.1 |
| *Cylindrospermopsis raciborskii* | GenBank: AF516730.1 |
| *Cylindrotheca closterium* | GenBank: GQ468542.1 |
| *Cymodocea nodosa* | GenBank: KT200607.1 |
| *Daucus carota* | GenBank: GQ380561.1 |
| *Deferribacter thermophilus* | NCBI Reference Sequence: NR_026043.1 |
| *Deinococcus geothermalis* | NCBI Reference Sequence: NR_074342.1 |
| *Deinococcus murrayi* | NCBI Reference Sequence: NR_026416.1 |
| *Desmarestia anceps* | GenBank: HE866895.1 |
| *Desmidium swartzii* | GenBank: AJ428133.1 |
| *Desulfitobacterium dehalogenans* | GenBank: L28946.1 |
| *Desulfobacter curvatus* | NCBI Reference Sequence: NR_041851.1 |
| *Desulfofaba gelida* | NCBI Reference Sequence: NR_028730.1 |
| *Desulfofrigus fragile* | NCBI Reference Sequence: NR_028732.1 |
| *Desulfofrigus oceanense* | NCBI Reference Sequence: NR_028731.1 |
| *Desulforhopalus vacuolatus* | NCBI Reference Sequence: NR_044653.1 |
| *Desulfotalea arctica* | NCBI Reference Sequence: NR_024949.1 |
| *Desulfotalea psychrophila* | NCBI Reference Sequence: NR_028729.1 |
| *Desulfotomaculum alkaliphilum* | NCBI Reference Sequence: NR_024947.1 |
| *Desulfotomaculum putei* | GenBank: HM228397.1 |
| *Desulfovibrio desulfuricans* | NCBI Reference Sequence: NR_036778.1 |
| *Desulfovibrio profundus* | NCBI Reference Sequence: NR_114641.1 |
| *Desulfovibrio salexigens* | NCBI Reference Sequence: NR_102801.1 |
| *Desulfurobacterium crinifex* | NCBI Reference Sequence: NR_114880.1 |
| *Desulfuromonas michiganensis* | GenBank: AF357915.2 |
| *Detonula confervacea* | GenBank: HQ912617.1 |
| *Diapensia lapponica* | GenBank: AF419794.1 |
| *Dinobryon divergens* | GenBank: EU025020.1 |
| *Ditylum brightwellii* | GenBank: X85386.2 |
| *Dunaliella tertiolecta* | GenBank: EF473747.1 |
| *Egeria densa* | GenBank: JF975484.1 |
| *Elodea canadensis* | GenBank: AF168841.1 |
| *Emiliania huxleyi* | GenBank: KC404141.1 |
| *Enterococcus faecalis* | GenBank: EU887827.1 |
| *Enteromorpha intestinalis* | GenBank: AJ000040.1 |
| *Erwinia amylovora* | GenBank: KM597069.1 |
| *Escherichia coli* | GenBank: AB269763.1 |
| *Eucalyptus globulus* | GenBank: HQ456544.1 |
| *Eucampia zodiacus* | GenBank: KC309495.1 |
| *Eucheuma isiforme* | GenBank: U25438.1 |
| *Ferroglobus placidus* | NCBI Reference Sequence: NR_074531.1 |
| *Ferroplasma acidarmanus* | GenBank: AF145441.1 |
| *Ferroplasma acidiphilum* | GenBank: AF513710.1 |
| *Ferroplasma cupricumulans* | GenBank: AY907888.1 |
| *Fervidobacterium pennavorans* | GenBank: EF565822.1 |
| *Fibrocapsa japonica* | GenBank: AY788931.1 |
| *Flavobacterium limicola* | GenBank: AB075232.1 |
| *Flexistipes sinusarabici* | NCBI Reference Sequence: NR_074881.1 |

Table S4 – *Continued from previous page*

| Species | Accession ID |
|---|---|
| *Fontinalis antipyretica* | GenBank: AF023714.1 |
| *Fragilaria barbararum* | GenBank: AJ971376.1 |
| *Fragilaria crotonensis* | GenBank: AM712616.1 |
| *Fragilariopsis cylindrus* | GenBank: EF140624.1 |
| *Fragilariopsis kerguelensis* | GenBank: KJ866919.1 |
| *Fucus gardneri* | GenBank: HQ710578.1 |
| *Gambierdiscus toxicus* | GenBank: EF202890.1 |
| *Gelidibacter gilvus* | NCBI Reference Sequence: NR_041693.1 |
| *Geobacillus caldoxylosilyticus* | GenBank: AY608951.1 |
| *Geobacillus stearothermophilus* | GenBank: EF025325.1 |
| *Geobacillus thermodenitrificans* | GenBank: AJ785764.1 |
| *Geobacillus thermoleovorans* | GenBank: JQ343209.1 |
| *Geoglobus ahangari* | NCBI Reference Sequence: NR_041788.1 |
| *Gephyrocapsa oceanica* | GenBank: KC404159.1 |
| *Gerbera jamesonii* | GenBank: AF107576.1 |
| *Glaciecola punicea* | NCBI Reference Sequence: NR_036866.1 |
| *Glycine max* | GenBank: X02623.1 |
| *Gonatozygon monotaenium* | GenBank: AJ428084.1 |
| *Gossypium hirsutum* | GenBank: L24145.1 |
| *Gracilaria verrucosa* | GenBank: M33638.1 |
| *Grammonema striatula* | GenBank: X77704.1 |
| *Guinardia flaccida* | GenBank: AJ535191.1 |
| *Gymnodinium breve* | GenBank: AF172714.1 |
| *Gymnodinium catenatum* | GenBank: AF022193.1 |
| *Gymnodinium mikimotoi* | GenBank: AF022195.1 |
| *Gymnodinium sanguineum* | GenBank: AJ415513.1 |
| *Gymnodinium veneficum* | GenBank: AF172712.1 |
| *Gyrodinium aureolum* | GenBank: AF172713.1 |
| *Gyrodinium instriatum* | GenBank: DQ084522.1 |
| *Haematococcus pluvialis* | GenBank: JQ315539.1 |
| *Haloanaerobium alcaliphilum* | GenBank: KU180221.1 |
| *Haloanaerobium lacusroseus* | NCBI Reference Sequence: NR_025924.1 |
| *Haloarcula vallismortis* | NCBI Reference Sequence: NR_116083.1 |
| *Halobacterium salinarum* | GenBank: AB663362.1 |
| *Halobaculum gomorrense* | GenBank: L37444.1 |
| *Halococcus morrhuae* | NCBI Reference Sequence: NR_043387.1 |
| *Haloferax volcanii* | NCBI Reference Sequence: NR_113448.1 |
| *Halogeometricum borinquense* | NCBI Reference Sequence: NR_028170.1 |
| *Halomonas campisalis* | GenBank: DQ077908.1 |
| *Halomonas elongata* | GenBank: KU053958.1 |
| *Halomonas marina* | GenBank: AJ306890.1 |
| *Halomonas subglaciescola* | NCBI Reference Sequence: NR_042067.1 |
| *Halonatronum saccharophilum* | NCBI Reference Sequence: NR_042717.1 |
| *Halorubrum saccharovorum* | NCBI Reference Sequence: NR_113484.1 |
| *Haloterrigena turkmenica* | NCBI Reference Sequence: NR_113515.1 |
| *Helianthus annuus* | GenBank: AF107577.1 |
| *Heliobacillus mobilis* | NCBI Reference Sequence: NR_040957.1 |
| *Heliobacterium modesticaldum* | NCBI Reference Sequence: NR_074517.1 |
| *Heterocapsa circularisquama* | GenBank: LC054932.1 |
| *Heterocapsa triquetra* | GenBank: AF022198.1 |
| *Heterosigma akashiwo* | GenBank: AB217869.1 |
| *Hordeum vulgare* | GenBank: AH001585.2 |

Table S4 – *Continued from previous page*

| Species | Accession ID |
|---|---|
| *Hormidium flaccidum* | GenBank: M95613.1 |
| *Hydrilla verticillata* | GenBank: KM982363.1 |
| *Hydrogenophaga pseudoflava* | NCBI Reference Sequence: NR_028717.1 |
| *Hydrogenophilus hirschii* | NCBI Reference Sequence: NR_104788.1 |
| *Ignicoccus hospitalis* | NCBI Reference Sequence: NR_028955.1 |
| *Ignicoccus islandicus* | NCBI Reference Sequence: NR_044910.1 |
| *Ignicoccus pacificus* | GenBank: AJ271794.1 |
| *Impatiens walleriana* | GenBank: L49285.1 |
| *Ipomoea batatas* | GenBank: HM053485.1 |
| *Isochrysis galbana* | GenBank: AJ246266.1 |
| *Isosphaera pallida* | NCBI Reference Sequence: NR_028892.1 |
| *Klebsiella oxytoca* | GenBank: AF390083.1 |
| *Klebsiella pneumoniae* | GenBank: KC990817.1 |
| *Koliella antarctica* | GenBank: AJ311569.1 |
| *Lactobacillus acidophilus* | GenBank: KC150145.1 |
| *Lactobacillus delbrueckii* | GenBank: KJ868760.1 |
| *Lactobacillus paracasei* | NCBI Reference Sequence: NR_121787.1 |
| *Lactobacillus rhamnosus* | NCBI Reference Sequence: NR_043408.1 |
| *Lactococcus lactis* | GenBank: KR604712.1 |
| *Lactococcus piscium* | GenBank: JN226414.1 |
| *Lactuca sativa* | GenBank: KT225377.1 |
| *Lantana camara* | GenBank: AJ236049.1 |
| *Larix decidua* | GenBank: AB026938.1 |
| *Larrea tridentata* | GenBank: AY929372.1 |
| *Lauderia annulata* | GenBank: DQ514849.1 |
| *Lemna minor* | GenBank: S67398.1 |
| *Lepidodinium chlorophorum* | GenBank: AB686253.1 |
| *Leptocylindrus danicus* | GenBank: AJ535175.1 |
| *Leptospirillum ferriphilum* | GenBank: AF356830.1 |
| *Leptospirillum ferrooxidans* | NCBI Reference Sequence: NR_074963.1 |
| *Limnothrix redekei* | GenBank: FM177493.1 |
| *Lingulodinium polyedrum* | GenBank: AB693195.1 |
| *Liriodendron tulipifera* | GenBank: AF206954.1 |
| *Listeria monocytogenes* | GenBank: M58822.1 |
| *Lithophyllum margaritae* | GenBank: KP192392.1 |
| *Lolium multiflorum* | GenBank: AY846367.1 |
| *Lolium perenne* | GenBank: AY519271.1 |
| *Marinithermus hydrothermalis* | NCBI Reference Sequence: NR_028639.1 |
| *Marinitoga piezophila* | NCBI Reference Sequence: NR_074102.1 |
| *Marinobacter alkaliphilus* | GenBank: EU440994.1 |
| *Mastigocladus laminosus* | GenBank: DQ431003.1 |
| *Merismopedia tenuissima* | GenBank: AJ639891.1 |
| *Mesotaenium kramstae* | GenBank: AJ553922.1 |
| *Methanobacterium subterraneum* | GenBank: JQ268007.1 |
| *Methanobacterium thermoaggregans* | GenBank: AF095264.1 |
| *Methanobacterium thermoautotrophicum* | GenBank: AF095262.1 |
| *Methanococcus jannaschii* | GenBank: M59126.1 |
| *Methanococcus thermolithotrophicus* | GenBank: M59128.1 |
| *Methanococcus voltae* | NCBI Reference Sequence: NR_074184.1 |
| *Methanococcus vulcanius* | NCBI Reference Sequence: NR_028701.1 |
| *Methanoculleus submarinus* | NCBI Reference Sequence: NR_028856.1 |
| *Methanogenium frigidum* | NCBI Reference Sequence: NR_104790.1 |

Table S4 – *Continued from previous page*

| Species | Accession ID |
|---|---|
| *Methanogenium frittonii* | GenBank: AJ862839.1 |
| *Methanohalophilus portucalensis* | GenBank: KT285318.1 |
| *Methanolobus psychrophilus* | GenBank: EF202842.1 |
| *Methanopyrus kandleri* | NCBI Reference Sequence: NR_074539.1 |
| *Methanosarcina barkeri* | GenBank: M59144.1 |
| *Methanothermobacter thermautotrophicus* | GenBank: DQ657903.1 |
| *Methanothermococcus okinawensis* | NCBI Reference Sequence: NR_028155.1 |
| *Methanothrix soehngenii* | NCBI Reference Sequence: NR_028242.1 |
| *Micrasterias americana* | GenBank: FR852595.1 |
| *Microcystis aeruginosa* | NCBI Reference Sequence: NR_074314.1 |
| *Microcystis wesenbergii* | GenBank: U40334.1 |
| *Moritella abyssi* | GenBank: AB554718.1 |
| *Moritella profunda* | NCBI Reference Sequence: NR_025381.1 |
| *Mucuna pruriens* | GenBank: AF525695.1 |
| *Mychonastes homosphaera* | GenBank: X73996.1 |
| *Nannochloropsis oceanica* | GenBank: FJ896231.1 |
| *Natrialba asiatica* | NCBI Reference Sequence: NR_113519.1 |
| *Natrinema pellirubrum* | NCBI Reference Sequence: NR_113528.1 |
| *Natronobacterium gregoryi* | NCBI Reference Sequence: NR_113531.1 |
| *Natronococcus occultus* | NCBI Reference Sequence: NR_113534.1 |
| *Natronomonas pharaonis* | NCBI Reference Sequence: NR_113497.1 |
| *Natronorubrum bangense* | NCBI Reference Sequence: NR_113538.1 |
| *Navicula arenaria* | GenBank: KJ961668.1 |
| *Navicula pelliculosa* | GenBank: AY485454.1 |
| *Nerium oleander* | GenBank: AF107572.1 |
| *Nicotiana tabacum* | GenBank: AJ236016.1 |
| *Nitrosotalea devanaterra* | GenBank: JN227488.1 |
| *Nitzschia dissipata* | GenBank: AJ867018.1 |
| *Nitzschia frigida* | GenBank: JQ582669.1 |
| *Nitzschia paleacea* | GenBank: AJ866996.1 |
| *Nitzschia sigma* | GenBank: AJ867279.1 |
| *Odontella aurita* | GenBank: HQ912687.1 |
| *Odontella mobiliensis* | GenBank: KC309500.1 |
| *Odontella regia* | GenBank: KC309502.1 |
| *Odontella sinensis* | GenBank: HQ912564.1 |
| *Olea europaea* | GenBank: L49289.1 |
| *Olisthodiscus luteus* | GenBank: AY788937.1 |
| *Oryza sativa* | GenBank: AF069218.1 |
| *Oscillatoria mougeotii* | GenBank: FJ434250.1 |
| *Ostreopsis ovata* | GenBank: AF244939.1 |
| *Palaeococcus helgesonii* | NCBI Reference Sequence: NR_029059.1 |
| *Pandorina morum* | GenBank: JQ315554.1 |
| *Papaver somniferum* | GenBank: DQ912867.1 |
| *Paracoccus halodenitrificans* | NCBI Reference Sequence: NR_025890.1 |
| *Paramecium tetraurelia* | GenBank: EF502045.1 |
| *Paraphysomonas imperforata* | GenBank: EF432519.1 |
| *Pavlova lutheri* | GenBank: AF102369.1 |
| *Pediastrum duplex* | GenBank: M62997.1 |
| *Pelagomonas calceolata* | GenBank: EF455763.1 |
| *Pelotomaculum thermopropionicum* | NCBI Reference Sequence: NR_040840.1 |
| *Peptostreptococcus productus* | NCBI Reference Sequence: NR_113270.1 |
| *Peridinium cinctum* | GenBank: AB185114.1 |

Table S4 – *Continued from previous page*

| **Species** | **Accession ID** |
|---|---|
| *Persephonella guaymasensis* | NCBI Reference Sequence: NR_025166.1 |
| *Persephonella marina* | NCBI Reference Sequence: NR_102828.1 |
| *Phaeocystis antarctica* | GenBank: JN381495.1 |
| *Phaeocystis globosa* | GenBank: AY851301.1 |
| *Phaeocystis pouchetii* | GenBank: AJ278036.1 |
| *Phaeodactylum tricornutum* | GenBank: GQ452861.1 |
| *Phyllogigas grandifolius* | GenBank: HE866931.1 |
| *Picea mariana* | GenBank: L01782.1 |
| *Picrophilus oshimae* | NCBI Reference Sequence: NR_026246.1 |
| *Pinus elliottii* | GenBank: AF051798.1 |
| *Pinus taeda* | GenBank: AH001728.2 |
| *Pisum sativum* | GenBank: U43011.1 |
| *Planktothrix agardhii* | GenBank: FJ159128.1 |
| *Planococcus halocryophilus* | GenBank: JF742665.1 |
| *Plantago lanceolata* | GenBank: AJ236046.1 |
| *Populus tremuloides* | GenBank: AF206999.1 |
| *Porphyra perforata* | GenBank: GU319856.1 |
| *Porphyra umbilicalis* | GenBank: AH010576.2 |
| *Porphyridium purpureum* | GenBank: AB045584.1 |
| *Posidonia australis* | GenBank: GQ497582.1 |
| *Posidonia oceanica* | GenBank: AY491942.1 |
| *Potamogeton perfoliatus* | GenBank: AY952389.1 |
| *Proboscia indica* | GenBank: AY485470.1 |
| *Prochlorococcus marinus* | NCBI Reference Sequence: NR_028762.1 |
| *Profundimonas piezophila* | NCBI Reference Sequence: NR_117943.1 |
| *Prorocentrum concavum* | GenBank: Y16237.1 |
| *Prorocentrum dentatum* | GenBank: AY551273.1 |
| *Prorocentrum gracile* | GenBank: AY443019.1 |
| *Prorocentrum lima* | GenBank: Y16235.1 |
| *Prorocentrum mexicanum* | GenBank: Y16232.1 |
| *Prorocentrum micans* | GenBank: AJ415519.1 |
| *Prorocentrum minimum* | GenBank: JF715165.1 |
| *Prunus persica* | GenBank: L28749.1 |
| *Prymnesium polylepis* | GenBank: AJ004866.1 |
| *Pseudoalteromonas antarctica* | NCBI Reference Sequence: NR_029317.1 |
| *Pseudoalteromonas haloplanktis* | GenBank: EU807989.1 |
| *Pseudochattonella verruculosa* | GenBank: AM075625.1 |
| *Pseudomonas aeruginosa* | GenBank: AM419153.2 |
| *Pseudomonas fluorescens* | GenBank: AY538263.1 |
| *Pseudomonas putida* | GenBank: KF278708.1 |
| *Pseudo-nitzschia fraudulenta* | GenBank: JN091721.1 |
| *Pseudo-nitzschia granii* | GenBank: GU373962.1 |
| *Pseudo-nitzschia multiseries* | GenBank: AM235382.1 |
| *Pseudo-nitzschia pseudodelicatissima* | GenBank: GU373965.1 |
| *Pseudo-nitzschia seriata* | GenBank: GU373969.1 |
| *Pseudoxanthomonas broegbernensis* | NCBI Reference Sequence: NR_025306.1 |
| *Pseudoxanthomonas taiwanensis* | NCBI Reference Sequence: NR_025198.1 |
| *Psychrobacter glacincola* | GenBank: AB334769.1 |
| *Psychrobacter muriicola* | NCBI Reference Sequence: NR_114669.1 |
| *Psychroflexus torquis* | GenBank: DQ007442.1 |
| *Psychromonas profunda* | NCBI Reference Sequence: NR_025506.1 |
| *Pyrobaculum aerophilum* | GenBank: L07510.1 |

Table S4 – *Continued from previous page*

| Species | Accession ID |
|---|---|
| *Pyrobaculum calidifontis* | NCBI Reference Sequence: NR_040922.1 |
| *Pyrobaculum islandicum* | NCBI Reference Sequence: NR_074372.1 |
| *Pyrobaculum oguniense* | NCBI Reference Sequence: NR_112094.1 |
| *Pyrobaculum organotrophum* | NCBI Reference Sequence: NR_112158.1 |
| *Pyrococcus abyssi* | NCBI Reference Sequence: NR_115145.1 |
| *Pyrococcus furiosus* | NCBI Reference Sequence: NR_074375.1 |
| *Pyrococcus glycovorans* | NCBI Reference Sequence: NR_029053.1 |
| *Pyrococcus horikoshii* | NCBI Reference Sequence: NR_115653.1 |
| *Pyrodinium bahamense* | GenBank: DQ500120.1 |
| *Pyrolobus fumarii* | NCBI Reference Sequence: NR_102985.1 |
| *Quercus rubra* | GenBank: AF132892.1 |
| *Quercus suber* | GenBank: GU476438.1 |
| *Ranunculus acris* | GenBank: AH001745.2 |
| *Rhizosolenia robusta* | GenBank: AY485481.1 |
| *Rhizosolenia setigera* | GenBank: AY485461.1 |
| *Rhodomonas salina* | GenBank: HM126532.1 |
| *Rosa hybrida* | GenBank: X66773.1 |
| *Roya anglica* | GenBank: AJ428081.1 |
| *Rubrobacter radiotolerans* | GenBank: U65647.1 |
| *Rubrobacter xylanophilus* | NCBI Reference Sequence: NR_074552.1 |
| *Ruppia maritima* | GenBank: JN034103.1 |
| *Salmonella enterica* | GenBank: KF535115.1 |
| *Scenedesmus acuminatus* | GenBank: AB037088.1 |
| *Scenedesmus acutus* | GenBank: AJ249512.1 |
| *Scenedesmus quadricauda* | GenBank: KC790429.1 |
| *Scrippsiella trochoidea* | GenBank: EF492513.1 |
| *Selenastrum minutum* | GenBank: AY846380.1 |
| *Serratia marcescens* | GenBank: GU991997.1 |
| *Setaria italica* | GenBank: KC996746.1 |
| *Shewanella gelidimarina* | GenBank: AY771753.1 |
| *Skeletonema ardens* | GenBank: DQ396522.1 |
| *Skeletonema costatum* | GenBank: JF489959.1 |
| *Skeletonema japonicum* | GenBank: DQ011160.1 |
| *Skeletonema marinoi* | GenBank: JF489953.1 |
| *Skeletonema menzelii* | GenBank: AJ535168.1 |
| *Skeletonema pseudocostatum* | GenBank: X85393.1 |
| *Skeletonema tropicum* | GenBank: EF138941.1 |
| *Solanum lycopersicum* | GenBank: KJ813729.1 |
| *Solanum tuberosum* | GenBank: FJ710157.1 |
| *Solenostemon scutellarioides* | GenBank: EU019244.1 |
| *Sorghum bicolor* | GenBank: AH001770.2 |
| *Sphaerospermopsis aphanizomenoides* | GenBank: GU197654.1 |
| *Sphagnum angustifolium* | GenBank: GQ375058.1 |
| *Sphagnum squarrosum* | GenBank: GQ375075.1 |
| *Spinacia oleracea* | GenBank: L24420.1 |
| *Spiroplasma apis* | NCBI Reference Sequence: NR_104858.1 |
| *Spiroplasma cantharicola* | NCBI Reference Sequence: NR_125516.1 |
| *Spiroplasma chinense* | NCBI Reference Sequence: NR_025698.1 |
| *Spiroplasma citri* | NCBI Reference Sequence: NR_036849.1 |
| *Spiroplasma clarkii* | NCBI Reference Sequence: NR_104750.1 |
| *Spiroplasma culicicola* | NCBI Reference Sequence: NR_025701.1 |
| *Spiroplasma diminutum* | NCBI Reference Sequence: NR_025702.1 |

Table S4 – *Continued from previous page*

| Species | Accession ID |
|---|---|
| *Spiroplasma floricola* | NCBI Reference Sequence: NR_025703.1 |
| *Spiroplasma insolitum* | NCBI Reference Sequence: NR_025705.1 |
| *Spiroplasma ixodetis* | NCBI Reference Sequence: NR_104852.1 |
| *Spiroplasma kunkelii* | NCBI Reference Sequence: NR_104847.1 |
| *Spiroplasma mirum* | NCBI Reference Sequence: NR_118707.1 |
| *Spiroplasma monobiae* | NCBI Reference Sequence: NR_104854.1 |
| *Spiroplasma sabaudiense* | NCBI Reference Sequence: NR_025710.1 |
| *Spiroplasma taiwanense* | NCBI Reference Sequence: NR_121701.1 |
| *Spiroplasma velocicrescens* | NCBI Reference Sequence: NR_025713.1 |
| *Spirulina platensis* | GenBank: AB074508.1 |
| *Staphylococcus aureus* | GenBank: DQ630753.1 |
| *Staphylococcus xylosus* | NCBI Reference Sequence: NR_036907.1 |
| *Staurastrum avicula* | GenBank: EF507555.1 |
| *Staurastrum pingue* | GenBank: AJ428109.1 |
| *Staurodesmus cuspidatus* | GenBank: EF507538.1 |
| *Stellarima microtrias* | GenBank: EU090011.1 |
| *Stephanodiscus hantzschii* | GenBank: DQ093370.1 |
| *Stephanopyxis palmeriana* | GenBank: AY485527.1 |
| *Streptococcus salivarius* | GenBank: M58839.1 |
| *Streptococcus thermophilus* | GenBank: AB812892.1 |
| *Stygiolobus azoricus* | NCBI Reference Sequence: NR_043434.1 |
| *Sulfobacillus benefaciens* | GenBank: EF679212.1 |
| *Sulfobacillus sibiricus* | NCBI Reference Sequence: NR_042730.1 |
| *Sulfobacillus thermosulfidooxidans* | GenBank: EU499919.1 |
| *Sulfobacillus thermotolerans* | GenBank: JX966410.1 |
| *Sulfolobus acidocaldarius* | NCBI Reference Sequence: NR_043400.1 |
| *Sulfolobus hakonensis* | NCBI Reference Sequence: NR_028222.1 |
| *Sulfolobus metallicus* | NCBI Reference Sequence: NR_043433.1 |
| *Sulfolobus tengchongensis* | NCBI Reference Sequence: NR_115150.1 |
| *Sulfolobus yangmingensis* | NCBI Reference Sequence: NR_028603.1 |
| *Sulfophobococcus zilligii* | NCBI Reference Sequence: NR_029316.1 |
| *Sulfurihydrogenibium kristjanssoni* | GenBank: AM778960.1 |
| *Sulfurisphaera ohwakuensis* | NCBI Reference Sequence: NR_043432.1 |
| *Symbiobacterium toebii* | GenBank: AF190460.1 |
| *Symbiodinium microadriaticum* | GenBank: KU900226.1 |
| *Synechococcus elongatus* | GenBank: HF678511.1 |
| *Synechococcus lividus* | GenBank: AF132772.1 |
| *Syntrophothermus lipocalidus* | NCBI Reference Sequence: NR_040796.1 |
| *Thalassionema nitzschioides* | GenBank: X77702.2 |
| *Thalassiosira allenii* | GenBank: HM991688.1 |
| *Thalassiosira constricta* | GenBank: KT692951.1 |
| *Thalassiosira curviseriata* | GenBank: AJ810859.1 |
| *Thalassiosira eccentrica* | GenBank: X85396.1 |
| *Thalassiosira guillardii* | GenBank: AF374478.2 |
| *Thalassiosira hendeyi* | GenBank: AM050629.1 |
| *Thalassiosira nordenskioeldii* | GenBank: DQ093365.1 |
| *Thalassiosira pseudonana* | GenBank: AY485452.1 |
| *Thalassiosira rotula* | GenBank: AF374480.2 |
| *Thalassiosira weissflogii* | GenBank: AY485445.1 |
| *Thermacetogenium phaeum* | NCBI Reference Sequence: NR_074723.1 |
| *Thermaerobacter nagasakiensis* | NCBI Reference Sequence: NR_024776.1 |
| *Thermoanaerobacter ethanolicus* | NCBI Reference Sequence: NR_044619.1 |

Table S4 – *Continued from previous page*

| Species | Accession ID |
|---|---|
| *Thermoanaerobacter keratinophilus* | NCBI Reference Sequence: NR_115188.1 |
| *Thermoanaerobacter mathranii* | GenBank: LC127101.1 |
| *Thermoanaerobacter siderophilus* | GenBank: KR736354.1 |
| *Thermoanaerobacter subterraneus* | GenBank: EU109461.1 |
| *Thermoanaerobacter tengcongensis* | GenBank: AF209708.1 |
| *Thermoanaerobacter yonseiensis* | GenBank: HM228410.1 |
| *Thermobrachium celere* | GenBank: DQ207958.2 |
| *Thermococcus alcaliphilus* | NCBI Reference Sequence: NR_040870.1 |
| *Thermococcus barophilus* | NCBI Reference Sequence: NR_042734.1 |
| *Thermococcus barossii* | NCBI Reference Sequence: NR_042735.1 |
| *Thermococcus celer* | NCBI Reference Sequence: NR_042736.1 |
| *Thermococcus chitonophagus* | NCBI Reference Sequence: NR_119236.1 |
| *Thermococcus fumicolans* | NCBI Reference Sequence: NR_042738.1 |
| *Thermococcus hydrothermalis* | NCBI Reference Sequence: NR_042740.1 |
| *Thermococcus peptonophilus* | NCBI Reference Sequence: NR_028193.1 |
| *Thermococcus siculi* | NCBI Reference Sequence: NR_028195.1 |
| *Thermocrinis ruber* | NCBI Reference Sequence: NR_121741.1 |
| *Thermodesulfovibrio yellowstonii* | NCBI Reference Sequence: NR_074345.1 |
| *Thermoplasma acidophila* | NCBI Reference Sequence: NR_028235.1 |
| *Thermoproteus uzoniensis* | NCBI Reference Sequence: NR_102955.1 |
| *Thermosipho japonicus* | NCBI Reference Sequence: NR_024726.1 |
| *Thermosphaera aggregans* | NCBI Reference Sequence: NR_074380.1 |
| *Thermosyntropha lipolytica* | NCBI Reference Sequence: NR_026356.1 |
| *Thermoterrabacterium ferrireducens* | GenBank: U76364.1 |
| *Thermotoga lettingae* | NCBI Reference Sequence: NR_074951.1 |
| *Thermotoga maritima* | NCBI Reference Sequence: NR_029163.1 |
| *Thermus aquaticus* | NCBI Reference Sequence: NR_025900.1 |
| *Thermus chliarophilus* | NCBI Reference Sequence: NR_026244.1 |
| *Thermus thermophilus* | NCBI Reference Sequence: NR_037066.1 |
| *Trichococcus patagoniensis* | GenBank: AF394926.1 |
| *Trichodesmium erythraeum* | NCBI Reference Sequence: NR_074275.1 |
| *Trifolium repens* | GenBank: AF071069.1 |
| *Triticum aestivum* | GenBank: AY049040.1 |
| *Tychonema bourrellyi* | GenBank: FJ184385.1 |
| *Ulva lactuca* | GenBank: KF419328.1 |
| *Vallisneria americana* | GenBank: AF069201.1 |
| *Veratrum californicum* | GenBank: AH003503.2 |
| *Vibrio marinus* | GenBank: AJ297540.1 |
| *Vitis vinifera* | GenBank: GQ849399.1 |
| *Volvox aureus* | GenBank: LC086362.1 |
| *Xanthomonas campestris* | GenBank: AF290420.1 |
| *Xylella fastidiosa* | NCBI Reference Sequence: NR_115924.1 |
| *Yersinia enterocolitica* | GenBank: M59292.1 |
| *Zea mays* | GenBank: AF168884.1 |
| *Zostera marina* | GenBank: HQ445940.1 |
| *Zostera noltii* | GenBank: AF207058.1 |

**Table S5.** Species names and Accession IDs of cbbL/rbcL gene sequences that were used in this study.

| Species | Accession ID |
|---|---|
| *Abies alba* | GenBank: AB029652.1 |
| *Abutilon theophrasti* | GenBank: HM849734.1 |

Table S5 – *Continued from previous page*

| Species | Accession ID |
|---|---|
| *Acer rubrum* | GenBank: DQ978428.1 |
| *Acidimicrobium ferrooxidans* | GenBank: GQ409765.1 |
| *Acidithiobacillus caldus* | GenBank: GQ409763.1 |
| *Acidithiobacillus ferrivorans* | GenBank: FJ467341.1 |
| *Acidithiobacillus ferrooxidans* | GenBank: GQ409767.1 |
| *Acidithiobacillus thiooxidans* | GenBank: GQ225727.1 |
| *Aldrovanda vesiculosa* | GenBank: AY096106.1 |
| *Amphiprora paludosa* | GenBank: FJ002140.1 |
| *Anabaena ucrainica* | GenBank: GU197741.1 |
| *Antirrhinum majus* | GenBank: GQ997015.1 |
| *Aphanizomenon ovalisporum* | GenBank: KP698018.1 |
| *Aplectrum hyemale* | GenBank: AF074108.1 |
| *Arabidopsis thaliana* | GenBank: KU739560.1 |
| *Arbutus unedo* | GenBank: KF997388.1 |
| *Aristotelia serrata* | GenBank: AF307904.1 |
| *Asterionella formosa* | GenBank: HQ912497.1 |
| *Asterionellopsis glacialis* | GenBank: HQ912510.1 |
| *Aulacoseira granulata* | GenBank: AB430659.1 |
| *Beta vulgaris* | GenBank: KM360669.1 |
| *Betula papyrifera* | GenBank: X56617.1 |
| *Betula pendula* | GenBank: KM360670.1 |
| *Brassica oleracea* | GenBank: GQ184376.1 |
| *Brassica rapa* | GenBank: KJ473492.1 |
| *Bryum argenteum* | GenBank: AY163024.1 |
| *Capsella bursa-pastoris* | GenBank: KT458036.1 |
| *Capsicum annuum* | GenBank: KJ773334.1 |
| *Carya glabra* | GenBank: L12637.2 |
| *Caulerpa serrulata* | GenBank: JQ745697.1 |
| *Ceratophyllum demersum* | GenBank: AB917052.1 |
| *Chamaebatiaria millefolium* | GenBank: U06797.1 |
| *Chamerion angustifolium* | GenBank: L10217.1 |
| *Chenopodium album* | GenBank: JX848451.1 |
| *Chlamydomonas reinhardtii* | GenBank: AB511846.1 |
| *Chlamydomonas subcaudata* | GenBank: GQ871929.1 |
| *Chlorella ellipsoidea* | GenBank: EU038287.1 |
| *Chlorella pyrenoidosa* | GenBank: EU038283.1 |
| *Chlorella saccharophila* | GenBank: AM260446.1 |
| *Chlorella sorokiniana* | GenBank: HM101339.1 |
| *Chlorella vulgaris* | GenBank: EU038286.1 |
| *Chondrus crispus* | GenBank: U02984.1 |
| *Chrysanthemum morifolium* | GenBank: KM218356.1 |
| *Cicer arietinum* | GenBank: AF308707.1 |
| *Citrus aurantium* | GenBank: AB505953.1 |
| *Citrus limon* | GenBank: AB505956.1 |
| *Closterium acerosum* | GenBank: AF203492.1 |
| *Coccolithus pelagicus* | GenBank: HQ656833.1 |
| *Coelastrum microporum* | GenBank: KP698027.1 |
| *Coscinodiscus concinnus* | GenBank: HQ912545.1 |
| *Coscinodiscus granii* | GenBank: HQ656838.1 |
| *Coscinodiscus jonesianus* | GenBank: KJ577887.1 |
| *Cosmarium biretum* | GenBank: AM911267.1 |
| *Cosmarium botrytis* | GenBank: AM911295.1 |

Table S5 – *Continued from previous page*

| **Species** | **Accession ID** |
|---|---|
| *Cosmarium crenatum* | GenBank: AM911268.1 |
| *Cosmarium meneghinii* | GenBank: AM911284.1 |
| *Cosmarium punctulatum* | GenBank: AM911289.1 |
| *Cosmarium subprotumidum* | GenBank: AM911292.1 |
| *Cryptomonas marssonii* | GenBank: AM051209.1 |
| *Cryptomonas ovata* | GenBank: AM051211.1 |
| *Cucumis sativus* | GenBank: L21937.1 |
| *Cyclotella cryptica* | GenBank: KM816805.1 |
| *Cyclotella meneghiniana* | GenBank: KM816803.1 |
| *Cylindrospermopsis raciborskii* | GenBank: JF895153.1 |
| *Cylindrotheca closterium* | GenBank: JX971010.1 |
| *Cymodocea nodosa* | GenBank: U80688.1 |
| *Daucus carota* | GenBank: KM360751.1 |
| *Desmarestia anceps* | GenBank: HE866816.1 |
| *Desmidium swartzii* | GenBank: HQ380525.1 |
| *Detonula confervacea* | GenBank: HQ912481.1 |
| *Diapensia lapponica* | GenBank: L12612.2 |
| *Ditylum brightwellii* | GenBank: DQ514766.1 |
| *Dunaliella tertiolecta* | GenBank: JQ039069.1 |
| *Egeria densa* | GenBank: AB004887.1 |
| *Elodea canadensis* | GenBank: DQ859167.1 |
| *Emiliania huxleyi* | GenBank: JX292160.1 |
| *Enteromorpha intestinalis* | GenBank: AF499671.1 |
| *Eucalyptus globulus* | GenBank: HM849985.1 |
| *Eucampia zodiacus* | GenBank: KC309568.1 |
| *Eucheuma isiforme* | GenBank: AF099691.1 |
| *Fibrocapsa japonica* | GenBank: AB280606.1 |
| *Fontinalis antipyretica* | GenBank: AB050949.1 |
| *Fragilaria crotonensis* | GenBank: HQ828187.2 |
| *Fragilariopsis cylindrus* | GenBank: EF423499.1 |
| *Fragilariopsis kerguelensis* | GenBank: KC920826.1 |
| *Gephyrocapsa oceanica* | GenBank: D45844.1 |
| *Gerbera jamesonii* | GenBank: L13643.1 |
| *Glycine max* | GenBank: Z95552.1 |
| *Gonatozygon monotaenium* | GenBank: FM992338.1 |
| *Gossypium hirsutum* | GenBank: JQ034248.1 |
| *Gracilaria verrucosa* | GenBank: JQ843364.1 |
| *Grammonema striatula* | GenBank: KF701600.1 |
| *Guinardia flaccida* | GenBank: KC309609.1 |
| *Gymnodinium breve* | GenBank: AY119786.1 |
| *Gymnodinium mikimotoi* | GenBank: JX899690.2 |
| *Haematococcus pluvialis* | GenBank: FJ438476.1 |
| *Helianthus annuus* | GenBank: L13929.1 |
| *Heterosigma akashiwo* | GenBank: HQ710629.1 |
| *Hordeum vulgare* | GenBank: LN626641.1 |
| *Hormidium flaccidum* | GenBank: EU477433.1 |
| *Hydrilla verticillata* | GenBank: KM982379.1 |
| *Impatiens walleriana* | GenBank: AB043508.1 |
| *Ipomoea batatas* | GenBank: JQ923431.1 |
| *Isochrysis galbana* | GenBank: HQ656829.1 |
| *Lactuca sativa* | GenBank: AY874437.1 |
| *Lantana camara* | GenBank: HM850104.1 |

Table S5 – *Continued from previous page*

| Species | Accession ID |
|---|---|
| *Larix decidua* | GenBank: FN689379.1 |
| *Larrea tridentata* | GenBank: Y15022.1 |
| *Lauderia annulata* | GenBank: DQ514769.1 |
| *Lemna minor* | GenBank: AM905730.1 |
| *Lepidodinium chlorophorum* | GenBank: AY331683.1 |
| *Leptocylindrus danicus* | GenBank: JX413575.1 |
| *Liriodendron tulipifera* | GenBank: AF190430.1 |
| *Lolium multiflorum* | GenBank: LT576830.1 |
| *Lolium perenne* | GenBank: AY395547.1 |
| *Mastigocladus laminosus* | GenBank: JQ918781.1 |
| *Mesotaenium kramstae* | GenBank: AJ553952.1 |
| *Microcystis aeruginosa* | GenBank: KP698052.1 |
| *Mucuna pruriens* | GenBank: EU128733.1 |
| *Mychonastes homosphaera* | GenBank: KC145515.1 |
| *Nannochloropsis oceanica* | GenBank: KT149178.1 |
| *Navicula pelliculosa* | GenBank: HQ337547.1 |
| *Nerium oleander* | GenBank: HM850198.1 |
| *Nicotiana tabacum* | GenBank: KC825342.1 |
| *Odontella aurita* | GenBank: HQ912551.1 |
| *Odontella mobiliensis* | GenBank: KC309574.1 |
| *Odontella regia* | GenBank: KC309576.1 |
| *Odontella sinensis* | GenBank: HQ912428.1 |
| *Olea europaea* | GenBank: DQ673304.1 |
| *Olisthodiscus luteus* | GenBank: AB280605.1 |
| *Oryza sativa* | GenBank: AJ746297.1 |
| *Pandorina morum* | GenBank: AB044165.1 |
| *Papaver somniferum* | GenBank: HM850232.1 |
| *Pavlova lutheri* | GenBank: HQ656830.1 |
| *Pediastrum duplex* | GenBank: EF078364.1 |
| *Pelagomonas calceolata* | GenBank: U89898.1 |
| *Phaeocystis antarctica* | GenBank: KP144261.1 |
| *Phaeocystis globosa* | GenBank: HQ656835.1 |
| *Phaeodactylum tricornutum* | GenBank: HQ912420.1 |
| *Phyllogigas grandifolius* | GenBank: HE866853.1 |
| *Picea mariana* | GenBank: EU364784.1 |
| *Pinus elliottii* | GenBank: AB081075.1 |
| *Pinus taeda* | GenBank: AF119177.1 |
| *Pisum sativum* | GenBank: JN661190.1 |
| *Planktothrix agardhii* | GenBank: EU151930.1 |
| *Plantago lanceolata* | GenBank: L36454.1 |
| *Populus tremuloides* | GenBank: AF206812.1 |
| *Porphyra perforata* | GenBank: AF452438.1 |
| *Porphyra umbilicalis* | GenBank: AF452446.1 |
| *Porphyridium purpureum* | GenBank: DQ308439.1 |
| *Posidonia australis* | GenBank: U80718.1 |
| *Posidonia oceanica* | GenBank: U80719.1 |
| *Potamogeton perfoliatus* | GenBank: AY952437.1 |
| *Proboscia indica* | GenBank: JQ315456.1 |
| *Prochlorococcus marinus* | GenBank: AY042090.1 |
| *Prunus persica* | GenBank: AF411493.1 |
| *Pseudochattonella verruculosa* | GenBank: AB280607.1 |
| *Pseudo-nitzschia fraudulenta* | GenBank: EF423503.1 |

Table S5 – *Continued from previous page*

| **Species** | **Accession ID** |
|---|---|
| *Pseudo-nitzschia granii* | GenBank: KU183494.1 |
| *Pseudo-nitzschia multiseries* | GenBank: KC801040.1 |
| *Pseudo-nitzschia pseudodelicatissima* | GenBank: KC801039.1 |
| *Quercus rubra* | GenBank: LM653089.1 |
| *Quercus suber* | GenBank: LM653092.1 |
| *Ranunculus acris* | GenBank: KF602170.1 |
| *Rhizosolenia robusta* | GenBank: JQ315467.1 |
| *Rhizosolenia setigera* | GenBank: HQ912425.1 |
| *Roya anglica* | GenBank: AJ553963.1 |
| *Ruppia maritima* | GenBank: HQ901576.1 |
| *Scenedesmus quadricauda* | GenBank: AB084332.1 |
| *Setaria italica* | GenBank: EF125138.1 |
| *Skeletonema ardens* | GenBank: JN162820.1 |
| *Skeletonema costatum* | GenBank: AF015569.1 |
| *Skeletonema japonicum* | GenBank: DQ514822.1 |
| *Skeletonema marinoi* | GenBank: KJ671816.1 |
| *Skeletonema menzellii* | GenBank: DQ514821.1 |
| *Skeletonema pseudocostatum* | GenBank: DQ514819.1 |
| *Skeletonema tropicum* | GenBank: KJ671817.1 |
| *Solanum lycopersicum* | GenBank: HF572813.1 |
| *Solanum tuberosum* | GenBank: KJ652187.1 |
| *Sorghum bicolor* | GenBank: AM849341.1 |
| *Sphaerospermopsis aphanizomenoides* | GenBank: FJ830541.1 |
| *Sphagnum angustifolium* | GenBank: AY309690.1 |
| *Sphagnum squarrosum* | GenBank: AY309706.1 |
| *Spirulina platensis* | GenBank: AY147205.1 |
| *Staurastrum pingue* | GenBank: AF203506.1 |
| *Stellarima microtrias* | GenBank: EU090032.1 |
| *Stephanodiscus hantzschii* | GenBank: AB831882.1 |
| *Stephanopyxis palmeriana* | GenBank: KP253080.1 |
| *Sulfobacillus thermosulfidooxidans* | GenBank: GQ409769.1 |
| *Thalassionema nitzschioides* | GenBank: KJ671820.1 |
| *Thalassiosira constricta* | GenBank: KT692950.1 |
| *Thalassiosira curviseriata* | GenBank: KJ671821.1 |
| *Thalassiosira eccentrica* | GenBank: DQ514789.1 |
| *Thalassiosira guillardii* | GenBank: DQ514796.1 |
| *Thalassiosira nordenskioeldii* | GenBank: KC985865.1 |
| *Thalassiosira pseudonana* | GenBank: HQ912419.1 |
| *Thalassiosira rotula* | GenBank: DQ514805.1 |
| *Thalassiosira weissflogii* | GenBank: DQ514811.1 |
| *Trichodesmium erythraeum* | GenBank: AB075924.1 |
| *Trifolium repens* | GenBank: KF602192.1 |
| *Triticum aestivum* | GenBank: LT576864.1 |
| *Ulva lactuca* | GenBank: EU484409.1 |
| *Vallisneria americana* | GenBank: U03726.1 |
| *Vitis vinifera* | GenBank: AJ635355.1 |
| *Volvox aureus* | GenBank: D63445.1 |
| *Zea mays* | GenBank: Z11973.1 |
| *Zostera marina* | GenBank: AB125348.1 |
| *Zostera noltii* | GenBank: U80733.1 |
